# Ubiquitous mRNA decay fragments in *E. coli* redefine the functional transcriptome

**DOI:** 10.1101/2021.11.18.469045

**Authors:** Lydia Herzel, Julian A. Stanley, Chun-Chen Yao, Gene-Wei Li

**Affiliations:** Department of Biology, Massachusetts Institute of Technology, Cambridge, MA 02142, USA; Harvard University, Cambridge, MA 02138, USA

**Keywords:** RNA sequencing, translation, mRNA decay

## Abstract

Bacterial mRNAs have short life cycles, in which transcription is rapidly followed by translation and degradation within seconds to minutes. The resulting diversity of mRNA molecules across different life-cycle stages impacts their functionality but has remained unresolved. Here we quantitatively map the 3′ status of cellular RNAs in *Escherichia coli* during steady-state growth and report a large fraction of molecules (median>60%) that are fragments of canonical full-length mRNAs. The majority of RNA fragments are decay intermediates, whereas nascent RNAs contribute to a smaller fraction. Despite the prevalence of decay intermediates in total cellular RNA, these intermediates are underrepresented in the pool of ribosome-associated transcripts and can thus distort quantifications for the abundance of full-length, functional mRNAs. The large heterogeneity within mRNA molecules *in vivo* highlights the importance in discerning functional transcripts and provides a lens for studying the dynamic life cycle of mRNAs.

## Introduction

Cellular mRNAs are present in different forms – nascent, full-length, partially degraded, and others. Whereas full-length molecules are often depicted as the canonical mRNAs in the central dogma of molecular biology, the other forms are necessary intermediates of their birth-death cycle and have potential to (mis)interact with the same pool of factors that regulate mRNA functionality, such as translation. A complete molecular census of the mRNA population is thus critical for quantifying the functional transcriptome, while also providing an opportunity to reveal the life cycle of mRNAs *in vivo*.

In bacteria, functional half-lives of mRNAs are often short, which could in principle give rise to a large heterogeneity among the transcripts that co-exist at the same time (Kuwano et al., 1977; Bernstein et al., 2002; Vargas-Blanco and Shell, 2020). After transcription initiates, mRNAs remain nascent for about a minute (considering a typical operon of 2 kilobases (kb) and a chain elongation rate of 45 nucleotides (nt)/s (Vogel et al., 1992; Proshkin et al., 2010)). Meanwhile, RNA endonucleases, such as RNase E in *E. coli*, typically cleave mRNAs within a few minutes (Cohen and McDowall, 1997; Hui et al., 2014; Mohanty and Kushner, 2016; Bechhofer and Deutscher, 2019; Trinquier et al., 2020). Following endonuclease cleavage, the upstream RNA fragments are digested by exonucleases, such as the 3′-to-5′ exonucleases PNPase and RNase II in *E. coli*, whereas the downstream fragments are further cleaved endonucleolytically (Donovan and Kushner, 1986). The timescales of these subsequent events are less characterized but can nevertheless determine the extent of decay intermediates present in the transcriptome (Fazal et al., 2015).

Amidst the heterogeneous RNA pool, translation takes place concurrently during transcription and mRNA decay in bacteria (Yarchuk et al., 1992). Whereas translation of nascent mRNAs is beneficial in *E. coli* by preventing Rho-dependent transcription termination, translation of mRNA decay intermediates may produce aberrantly truncated proteins and require rescue of ribosomes that are stalled at the end of RNA fragments (Richardson, 2002; Keiler, 2015; Müller et al., 2021). In the meantime, some mRNA decay intermediates are translation-incompetent but often included by RNA quantification methods as the total cellular RNA (Laalami et al., 2014). These considerations highlight the need to quantitatively resolve the forms of mRNA molecules across the transcriptome.

Existing methodologies for quantification primarily rely on regional information and are thus blind to distinct forms of RNAs. RT-qPCR and conventional RNA-seq detect ∼100 bases at a time. Although Northern blotting could distinguish RNAs of different lengths, it only detects transcripts containing the short stretch of complementary sequence that is probed. Further, partial RNA fragments in Northern blots are often spread out over a large size range that dilutes the signal compared to full-length RNAs. Recently, several high-throughput single-molecule strategies have been employed to explore major RNA isoforms (SMRT-Cappable-seq, SEnd-seq, Oxford Nanopore technologies) (Yan et al., 2018; Ju et al., 2019; Grünberger et al., 2021), which could in principle be used to probe the presence of nascent or partially degraded RNAs. However, these methods involve at least one size-selection step that restricts the range of RNAs (or cDNAs) analyzed. Therefore, the full spectrum of RNA molecules present in the cell remains unmapped.

Here we present a molecular census of RNA 3′ ends across the *E. coli* transcriptome. The 3′ terminal position of RNAs varies during their life cycle, and the relative representations can be quantified using methods that are not necessarily restricted to a limited size range (Mondal et al., 2016; Dar and Sorek, 2018). We find that, during steady-state growth, the majority of RNA molecules do not possess the mature 3′ ends of the corresponding transcription units (TUs). Instead, the 3′ termini are widely distributed across positions internal to TUs. Most RNA fragments at steady state are decay intermediates and not nascently transcribed. By contrast, translation primarily takes place on full-length and nascent RNAs. Together, our results indicate that the *E. coli* transcriptome consists of substantial amounts of both functional and non-functional mRNAs, which are often indiscernible by existing quantification methods.

## Results

### A molecular census of *in vivo* mRNA states by 3′ end sequencing

To obtain a comprehensive molecular census of mRNAs *in vivo*, the methods of RNA extraction and detection must be able to capture every type of molecule with a similar efficiency. They must also minimize additional RNA fragmentation during and after cell lysis. Here we describe our approach to meet these criteria.

To detect every type of RNA molecule throughout its life cycle, we take advantage of the ability of RNA 3′ end ligation to quantitatively mark each molecule containing a 3′ hydroxyl group with a preadenylated DNA adapter (Fig. 1A)(Churchman and Weissman, 2011; Carrillo Oesterreich et al., 2016). Deep sequencing of ligation junctions can be carried out to determine the 3′ end position, henceforth referred to as 3′ end sequencing (conceptually similar to previously described approaches (Dar et al., 2016; Mondal et al., 2016)). Compared to 5′ end positions, sequences at RNA 3′ ends allow us to distinguish transcripts with mature TU ends from those that are nascent or decay intermediates. Although the exact nature of RNA length remains undetermined, 3′ end sequencing provide a partial molecular census without excluding certain size ranges that is currently a component of existing full-length mapping strategies (Yan et al., 2018; Ju et al., 2019; Grünberger et al., 2021).

**Figure 1:**
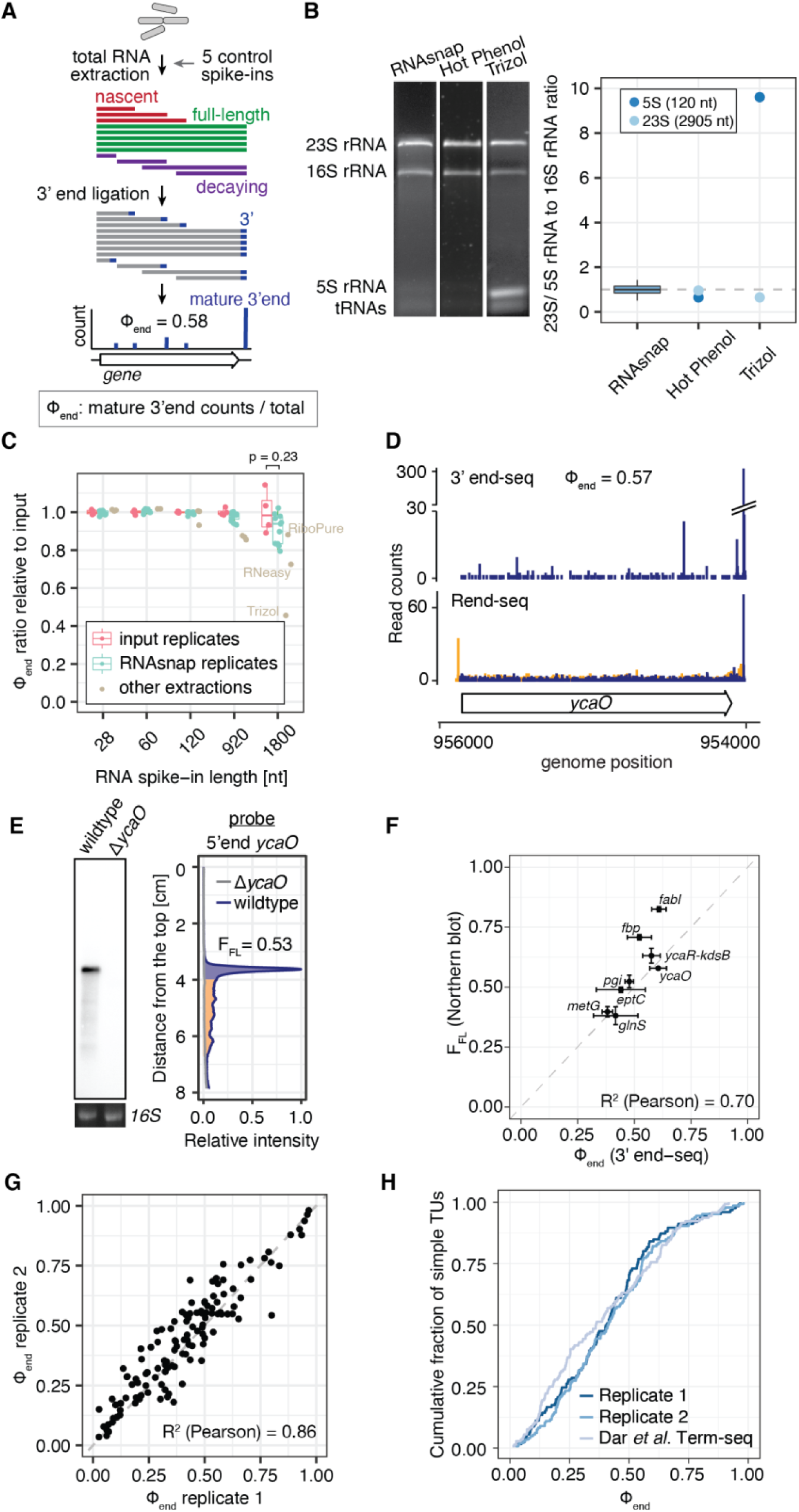
*In vivo* RNA 3′ end profiles reveal that the majority of transcripts are not full-length in *E. coli*. (A) Schematic of 3′ end sequencing and hypothetical data derived from the suite of nascent, decaying and full-length transcripts. The measure Φ_end_ is calculated as the fraction of 3′ end sequencing reads that map to the 3′ end of the mature transcript. Φ_end_ represents an upper bound for the fraction of full-length mRNA. (B) Minimization of length bias in RNA extraction. Left: Representative total RNA profiles by agarose gel electrophoresis from 3 different RNA extraction methods. Right: Molar ratio of rRNAs measured by 3′ end-seq. The mean 5S or 23S to 16S rRNA 3′ end ratio ± standard deviation for RNAsnap samples is 1.02 ± 0.22. The dashed line indicates the expected ratio of 1. Boxplot whiskers indicate 1.5 times the interquartile range. (C) Integrity of spiked-in reference RNAs. Five reference RNAs are synthesized *in vitro* (‘input’) and spiked in during RNA extraction. Boxplot shows Φ_end_ of the reference RNAs (spiked-in or input) normalized by the mean Φ_end_ of the input. Several technical replicates are shown for RNAsnap and input. The p-value was calculated between “input” and “RNAsnap” replicates with the Wilcoxon rank-sum test. No statistical test was applied to compare “other extractions” to “input” as this is a heterogenous group of samples from multiple RNA extractions with different characteristics. (D) 3′ end- and Rend-seq data for the simple transcription unit (TU) encoding *ycaO*. For 3′ end-seq, 3′-mapped (blue) read counts are plotted. Each read corresponds to one RNA 3′ end. For Rend-seq, 5′-mapped (orange) and 3′-mapped (blue) read counts are plotted separately to illustrate transcript boundaries. Here, a 5′ or 3′-mapped read corresponds to one RNA fragments generated during library preparation. (E) Northern blot (left) and intensity profile (right) for the same TU as in D. F_FL_ is calculated as the ratio of full-length signal intensity (blue) over the total intensity (orange + blue). The probe hybridizes to the 5′ region of the TU. 16S rRNA detected with SybrSafe staining after agarose gel electrophoresis prior transfer is shown as loading control. (F) Correlation between 3′ end sequencing and Northern blot analysis. Φ_end_ and F_FL_ for 8 simple TUs, ranging from 0.9-2.1kb, are plotted. Error bars reflect standard deviation of 3 and 2 replicates from Northern blotting and 3′ end-seq, respectively. Full northern blots are given in Fig. S1A. (G) Reproducibility of Φ_end_ by 3′ end sequencing of RNAsnap-extracted RNA. Scatter plot shows Φ_end_ from simple TUs between two biological replicates. (H) Cumulative distributions of Φ_end_ for two biological replicates and published data from (Dar and Sorek, 2018), with medians around 0.4.

To minimize biases and fragmentation introduced during RNA extraction, we tested several extraction methods and found RNAsnap to have the least amount of impact (Stead et al., 2012). Several metrics were used to evaluate the quality of extracted RNAs. First, the three ribosomal RNAs (5S, 16S, and 23S) spanning a size range of 120 nt to 2905 nt are expected to be in similar molar concentrations (Bremer and Dennis, 1996; Dong et al., 1996). Using 3′ end sequencing, we found that both RNAsnap and hot-phenol extraction reproduce the expected molar ratios among rRNAs (1.02 ± 0.22, mean ± standard deviation), whereas the commonly used TRIzol reagent strongly favors the shorter 5S rRNA as reported earlier (Fig. 1B)(Stead et al., 2012). Second, we spiked in *in vitro* synthesized (chemically synthesized or transcribed) reference RNAs during extraction to examine the degree of fragmentation after cell harvesting. As a measure for the integrity of each RNA species, we define Φ_end_ as the fraction of the corresponding 3′ end sequencing reads that map to the full-length 3′ end. We found that Φ_end_ for RNAsnap-extracted sample remains largely unchanged across different sizes of reference RNAs (Fig. 1C). By contrast, other extraction methods yielded reduced Φ_end_ for longer reference RNAs, indicating damages or skewed size representation during extraction. Third, additional evidence based on Northern blotting and density fractionation, as presented below, confirms that the RNA molecules observed using 3′ end sequencing of RNAsnap-extracted samples are faithful representations of the heterogeneity *in vivo*.

### Full-length transcripts constitute a small molar fraction of total mRNA

The robust extraction and mapping methods enable us to examine the diverse RNA forms present in the *E. coli* transcriptome during exponential growth. As expected, the positions that correspond to mature 3′ ends of a TU often have the most prominent signals (Fig. 1D). However, a substantial amount of RNA 3′ ends are also observed at many positions internal to the TU, suggesting a large transcriptome heterogeneity. To quantify this heterogeneity, we again use Φ_end_ to denote the fraction of reads that map to a transcript’s mature 3′ end for each transcription unit (TU). Because the RNAs that end at mature 3′ ends include both full-length and some decay intermediates (Fig. 1A), Φ_end_ represents an upper bound for the fraction of RNA molecules that are full-length. For a representative transcript encoding *ycaO*, the molar fraction of full-length RNAs is less than 57% (Φ_end_ = 0.57, Fig. 1D). Overall, the median Φ_end_ across all simple TUs (defined as those with a single promoter and a single terminator, Methods) is 0.4 and highly reproducible (Fig. 1G-H), indicating that full-length transcripts are a minority among total mRNA molecules.

Confirming this result, Northern blot analysis against the 5′ portion of simple TUs detected both full-length transcripts and partial ones of diverse lengths (Fig. 1E and S1A). Similar to 3′ end sequencing results, Northern blot signals from partial transcripts are dimly scattered across a wide swath, but nevertheless add up to a substantial fraction of total signals. Northern blots provide an estimate for the fraction of probe-binding RNAs that are full-length (F_FL_). Although neither Φ_end_ or F_FL_ represents the exact fraction of full-length RNAs, they showed good correspondence among the 8 simple TUs that we tested (Fig. 1F). These results indicate that the low Φ_end_ is not an artifact of the sequencing method. Additional confirmatory evidence comes from previously published transcriptomic data in *E. coli*. Term-seq is a similar method to 3′ end sequencing but uses a different RNA extraction kit and library design (Dar and Sorek, 2018). Despite these differences, it showed a similar distribution of Φ_end_ as our data (Fig. 1H, S1B). Overall, we conclude that the molar fraction of full-length RNA is small.

### Majority of partial transcripts are not nascent

Next, we quantified the contribution of nascent transcripts to the surprisingly high fraction of internal 3′ ends. If most partial transcripts were nascent, we would expect internal 3′ ends from total RNA and nascent RNA to follow similar patterns of RNA polymerase (RNAP) pausing across the body of TUs. Nascent transcripts are associated with RNAP as a part of the biochemically stable transcription elongation complex and can be purified by immunoprecipitation (Larson et al., 2014; Vvedenskaya et al., 2014) (Fig. 2A, S2, Methods). Sequencing data for 3′ ends of RNAP-associated transcripts showed strikingly different profiles from the 3′ ends of total RNAs (Fig. 2B). Furthermore, the elemental pause sequence for RNAP was recapitulated in nascent RNA but not in total RNA (Imashimizu et al., 2015; Larson et al., 2014; Vvedenskaya et al., 2014) (Fig. 2C). These two observations indicate that most partial transcripts were not nascent.

**Figure 2:**
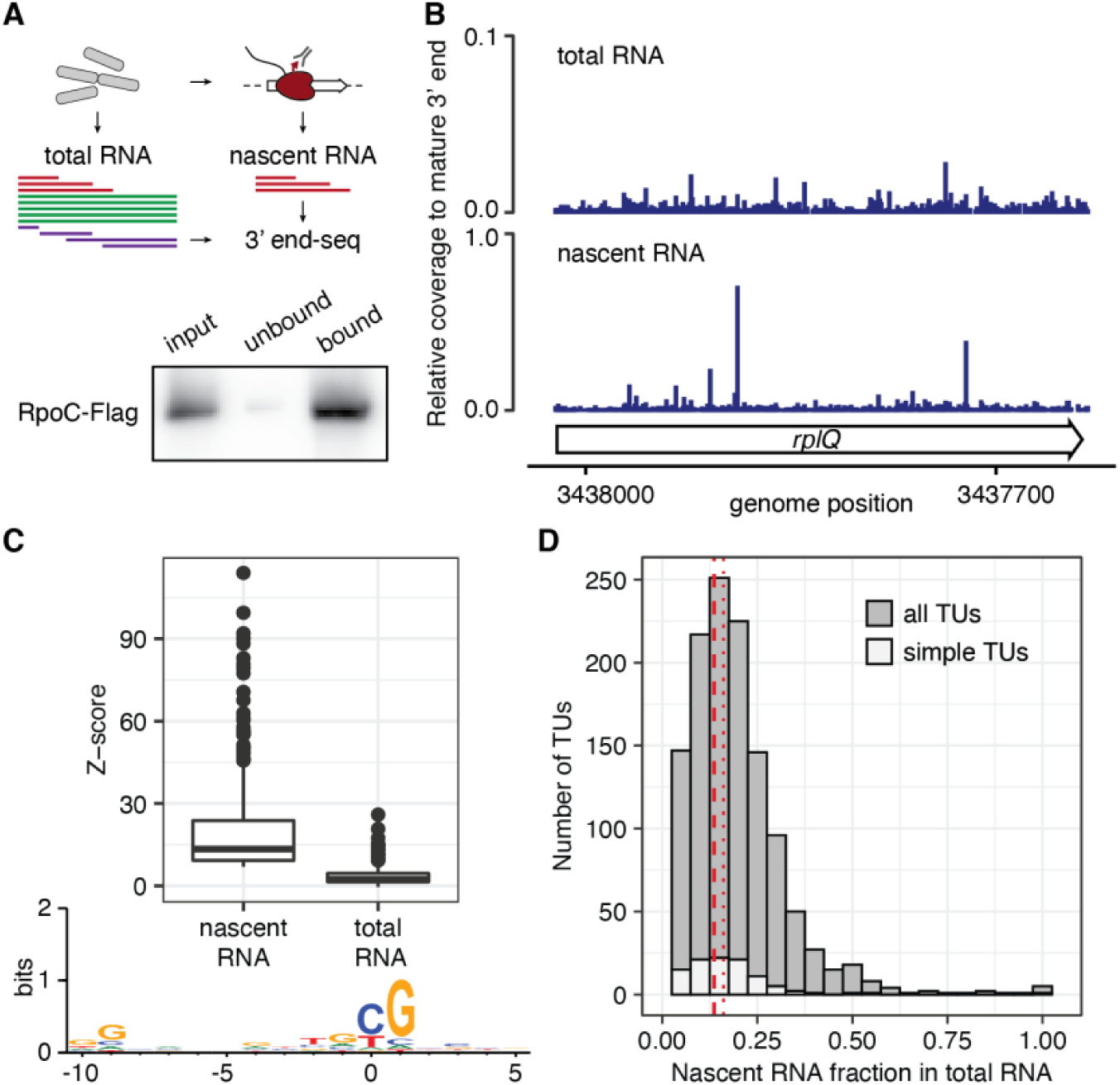
Nascent RNAs contribute to a small fraction of the transcriptome. (A) Schematic of 3′ end sequencing of nascent and total RNA. Nascent RNA is enriched by immunoprecipitation of Flag-tagged RNA polymerase (RpoC-Flag) and subject to 3′ end sequencing. Lower panel shows efficient isolation of RNA polymerases by western blot analysis with the Flag-antibody detecting RpoC-FLAG prominently in the total lysate (input) and eluate (bound), but less in the unbound fraction after immunoprecipitation. Full western in Fig. S2. (B) 3′ end sequencing results for nascent and total RNA. Internal 3′ end read counts are normalized to the read count at the mature 3′ end of *rplQ*. (C) Signals at transcriptional pause sites. Transcriptional pause sites are identified as peaks in 3′ end signals for nascent RNAs. Boxplot compares Z-score distributions for these sites in nascent RNA and total RNA. Sequence logo for regions around transcriptional pause sites resembles the elemental pause sequence. (D) Estimates of nascent RNA fraction in total RNA. Histogram shows the distribution of estimated fraction of total RNA that is nascent across different TUs. Simple TUs are a subset of all TUs and overlayed as white bars. Red lines mark distributions medians (dashed – simple TU, dotted – all TUs).

To estimate the nascent RNA fraction for each TU in total RNA, we compared the fraction of 3′ ends that map to elemental pause sites, denoted as f_pause_, in both the total and nascent RNA samples. If pause site signals in total RNAs are entirely contributed by nascent RNAs, the nascent RNA fraction in total RNAs can be calculated as the ratio of f_pause_ between the total and nascent RNA samples. In reality, pause site signals in total RNAs may be also derived from decaying RNAs. Therefore, the ratio of f_pause_s provides an upper bound for nascent transcripts in the total RNA pool. The distribution median was 0.15, indicating that nascent RNAs contribute at most 15% of all RNA molecules for a typical transcription unit (Fig. 2D).

### Majority of partial transcripts are decay intermediates

If a small fraction of partial transcripts is nascent, the remainder is possibly partially degraded. In this model, RNAs with longer half-lives should correspond to a low prevalence of RNA decay intermediates and thus a high Φ_end_. Indeed, stable noncoding RNAs showed Φ_end_ close to 1, whereas messenger RNAs of similar lengths have much lower Φ_end_ (Fig. 3A). Among mRNAs, Φ_end_ also correlates positively with mRNA half-life (Fig. S3A, R (Pearson) = 0.46). Because genome-wide measurements of RNA half-lives do not report full-length RNA stability (see below), we do not expect a perfect correlation between half-lives and Φ_end_. These results suggest that most RNA molecules bearing internal 3′ ends could be generated during the degradation process.

**Figure 3:**
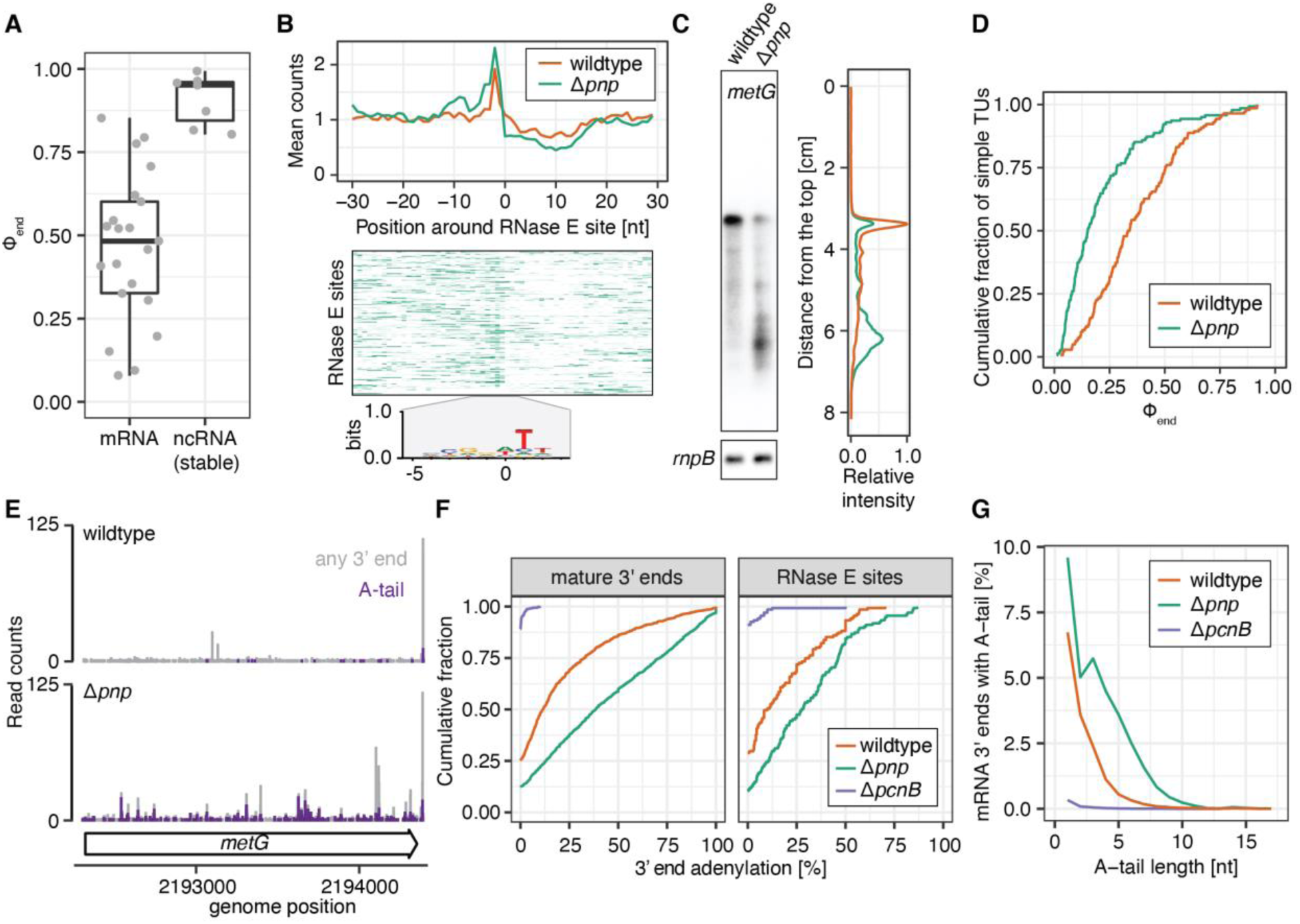
*In vivo* RNA decay signatures corroborate high amounts of decaying transcripts. (A) Distribution of Φ_end_ for size-matched mRNAs and stable noncoding RNAs (ncRNA). Boxplot shows Φ_end_ for stable ncRNAs (*ffs, gcvB, rnpB*, 5S, 16S, 23S rRNAs, *ssrA*) and mRNAs whose full-length products are in the same size range as the former. Whiskers correspond to 1.5 times the interquartile range. (B) Enrichment of 3′ end signal at putative RNase E cleavage sites. Top: Average 3′ end signals surrounding 1423 putative RNase E cleavage sites that has at least 30 reads in a 60-nt window. For each site, the read count for each position is normalized to the average read count in a symmetric 60-nt window. Line plots show the mean normalized count at each position across all sites. Middle: Normalized 3′ end read counts across individual RNase E sites in for Δ*pnp*. Each row represents data from each putative RNase E site. Color intensity represents the normalized 3′ end signal across the 60-nt window. Rows are sorted by descending Z-score at the putative cleavage site. The 200 sites with the highest number of reads in the 60-nt window are shown. (See corresponding wildtype heatmap in Fig. S3B and full heatmaps in Fig. S3C) Bottom: Sequence logo at these putative RNase E cleavage sites. (C) Accumulation of partial transcripts in Δ*pnp*. Left: Northern blot probing the 5′ region of *metG* in wildtype and Δ*pnp* shows decreased RNA integrity in cells lacking PNPase. Probe against stable ncRNA *rnpB* as loading control. Right: Intensity profiles of the Northern blot. See analyses for more TUs and replicates in Fig. S1A. (D) Global decrease in full-length RNAs in the *pnp* deletion (Δ*pnp*) compared to wildtype. Cumulative distributions of Φ_end_ for Δ*pnp* and wildtype are plotted for simple TUs. (E-G) Prevalence of short PolyA polymerase (PcnB)-dependent polyA tails. (E) 3′ end profiles across *metG* in wildtype and Δ*pnp*. Coverage shown in purple corresponds to adenylated 3′ ends, grey coverage corresponds to all 3′ ends. (F) Cumulative distributions of the percent of adenylation for mature 3′ ends and putative RNase E sites (positions with at least 10 reads). Adenylation is abolished upon deletion of the PolyA polymerase (*ΔpcnB*) and increases upon deletion of the exonuclease PNPase (*Δpnp*). (G) Percent of mRNA 3′ ends with A-tails of different lengths (includes mature and internal 3′ ends). The average A-tail length is very short and increases upon deletion of the exonuclease PNPase (*Δpnp*).

We next examined the location of internal 3′ ends and its relationship to RNA-decay pathways. In *E. coli*, mRNA degradation primarily occurs through an initial endonuclease cleavage, followed by 3′-5′ exonucleolytic activities and further rounds of endonuclease cleavage. RNase E is the main endonuclease and is essential for viability. PNPase is the main RNA exonuclease whose function can be partly compensated by other 3′-5′ exonucleases, such as RNase II (Hui et al., 2014; Mohanty and Kushner, 2016). To test if RNase E cleavages could explain the internal 3′ end profiles, we compared our wildtype 3′ end-seq data to previously published 5′-end sequencing data for an RNase E deficient *E. coli* strain (Clarke et al., 2014). We found that 3′ end signal is enriched among 1423 putative RNase E cleavage sites that have sufficient read coverage in our data, and the enrichment is even more pronounced in the strain lacking PNPase (Fig. 3B, S3B-C). Consistently, partial transcripts accumulate upon PNPase deletion, as indicated by both 3′ end sequencing and Northern blots (Fig. 3C-D). These results suggest that endonucleolytic cleavage followed by exonucleolytic digestion contributes to the ubiquitous internal 3′ ends that we observed.

Further supporting the prevalence of decay intermediates *in vivo*, we found signatures of adenylated 3′ ends (Fig. 3E). RNA 3′ end adenylation has been identified as a decay enhancing posttranscriptional RNA modification in *E. coli* (Kushner, 2015). Adenylation is mediated by the polyA polymerase PcnB and facilitates 3′-5′ exonucleolytic decay, especially for structured RNA ends. Indeed, our 3′ end sequencing data show a substantial level of adenylation at mature TU ends (median = 11.5% in wildtype, 38.5% in Δ*pnp*, 0.0% in Δ*pcnB*, Fig. 3F). Internal 3′ ends that map to putative RNase E cleavage sites also show adenylation in a manner that depends on PNPase and PcnB (median = 10.0% in wildtype, 27.8% in Δ*pnp*, 0.0% in Δ*pcnB*, Fig. 3F, S3D). These adenylated internal 3′ ends likely represent transient intermediates between actions of the polyA polymerase and 3′-5′ exonucleases. Overall, the length of A-tails responds similarly to deletions of PNPase and PcnB: The median A-tail length is short with 2 nt in wildtype cells, consistent with what was described previously (Waters et al., 2017). It increases to 3 nt in cells lacking PNPase, and vanishes in cells lacking PcnB (Fig. 3G, S3). Together, these lines of evidence support the model that many partial transcripts are RNA decay intermediates.

Interestingly, we found a higher overall Φ_end_ in a different bacterial species, *Bacillus subtilis*, that possesses an additional degradation pathway via the 5′-3′ exonuclease RNase J (Mathy et al., 2007). The median Φ_end_ is 0.6 for *B. subtilis*, compared to 0.4 for *E. coli* under a similar growth condition (Fig. S3I-K). This difference is consistent with the facts that mRNA decay in *B. subtilis* may not require sequential rounds of endonuclease cleavage, and that transcription elongation is much faster in *B. subtilis* (Hui et al., 2014; Johnson et al., 2020).

### Translation primarily occurs on a small fraction of transcripts

Among the majority of mRNA molecules that are decay intermediates, many may lack ribosome binding sites and hence are incompetent for translation. Indeed, density fractionation by sucrose gradients shows that partial transcripts are depleted in ribosome-rich fractions, as indicated by increased Φ_end_ in heavier fractions (Fig. 4A). The increased Φ_end_ beyond the input levels further confirms that the limited Φ_end_ observed in total RNA is not a result of post-lysis fragmentation and that partial transcripts contribute to a smaller faction of translated RNAs. Consistent with the expectation that decay intermediates are incompetent for translation, we found that RNAs that end at putative RNase E cleavage sites are much more prevalent in the ribosome-free fraction than the ribosome-rich fractions (Fig. 4B).

**Figure 4:**
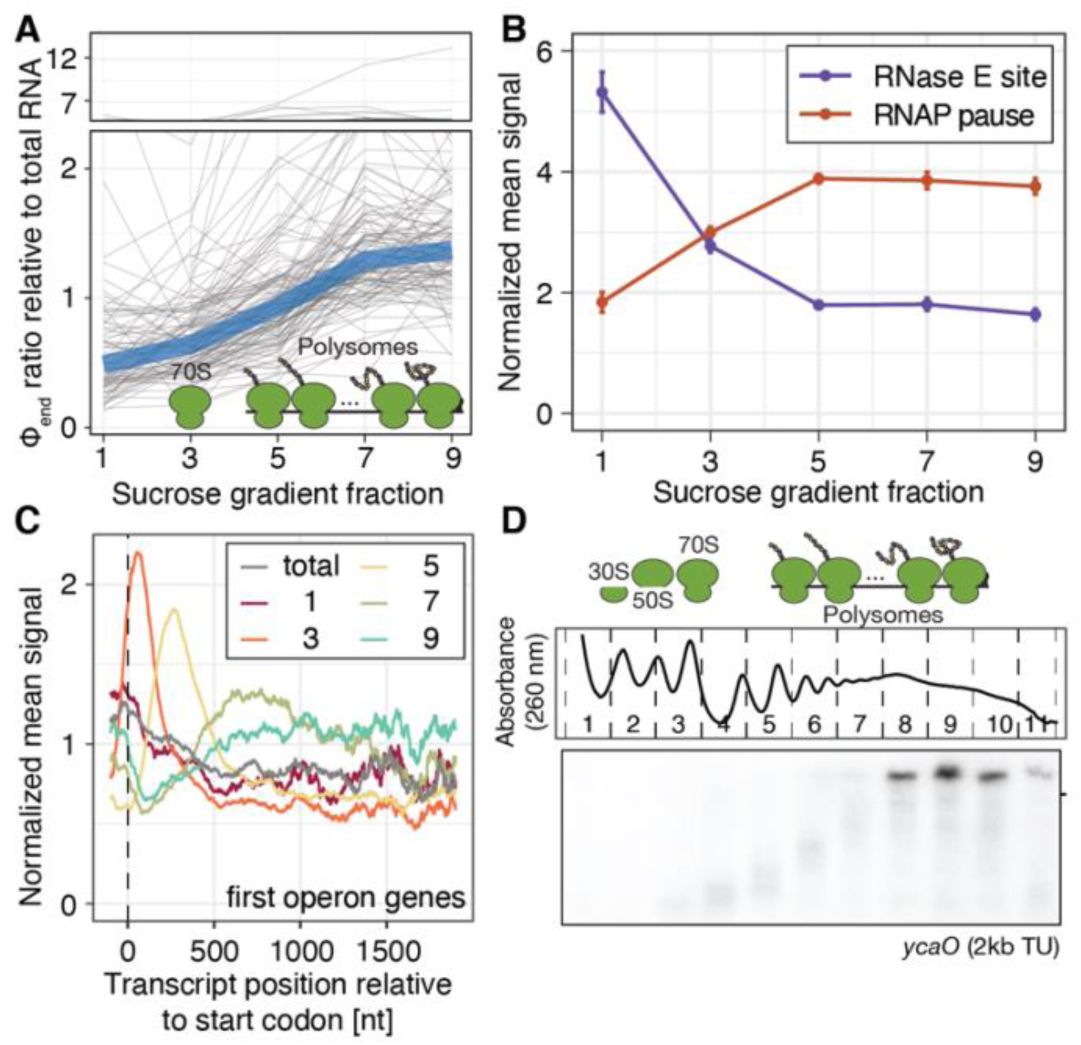
Enrichment of translation-competent RNAs in polysomes. (A) Global increase of Φ_end_ in heavy fractions of sucrose gradient. For each simple TU, the Φ_end_ in each fraction relative to the input Φ_end_ is plotted in grey. The median among all simple TUs is shown in blue, which starts below the input in lighter fractions and rises above the input in heavier fractions. The ribosome cartoon illustrates where monosomes (fraction 3) and polysomes (fractions >3) sediment across the gradient (compare absorbance profile in D). (B) Signatures of nascent RNA and decay intermediates across sucrose gradient. For each fraction, 3′ end signal at each pause site (or putative RNase E cleavage site) is first normalized to the mean signal in a symmetrical 60 nt window, and then averaged across 396 RNA polymerase pause sites (or 460 putative RNase E sites). Error bars reflect standard error of the mean. (C) Enrichment of 3′ end positions relative to start codons in different polysome fractions. For 538 first CDSs in operons, the average 3′ end signal normalized to the CDS mean signal (Methods) is plotted with respect to distance to the start codon for individual fractions. (D) Size distribution for partial mRNAs in different polysome fractions. Top: Absorbance profile across the sucrose gradient. Fraction numbers and the peaks that associate with 30S, 50S, 70S and polysomes are labeled. Bottom: Northern blot analysis probing the 5′ region of *ycaO* across different fractions.

We expect that translated partial mRNAs primarily consist of nascent mRNAs. Indeed, unlike total RNAs, RNAP pause site signals are prevalent in ribosome-rich fractions (Fig. 4B). Further, the 3′ ends of the monosome fraction are enriched immediately downstream of the first start codons in operons, whereas 3′ ends of higher polysome fractions are enriched at increasing distances from start codons (Fig. 4C). Northern blot analysis confirms that partial transcripts increase in size with increasing sucrose density, giving way to full-length transcripts at heaviest density (Fig. 4D, S4). Together, these results confirm that, despite a high molar fraction of decaying transcripts in the transcriptome, most translation occurs on nascent and full-length transcripts.

### Quantitation of the functional transcriptome requires discerning decay intermediates

The prevalence of translation-incompetent mRNAs in the transcriptome suggests that many RNA quantification methods may not accurately measure the levels of functional mRNAs. To demonstrate this potential problem, we compared *E. coli* RNA-seq data for wildtype cells and cells lacking PNPase. Previous studies using DNA microarray or RNA-seq, which does not distinguish functional and partial RNAs, have noted limited changes in RNA abundance and half-lives in the absence of PNPase (Bernstein et al., 2004; Pobre and Arraiano, 2015). This result was recapitulated in data from our own lab (Fig. 5A)(Lalanne et al., 2018). However, 3′ end sequencing showed that the full-length RNA fractions are dramatically reduced, as shown earlier in Fig. 3D. For example, the gene *fbp* has similar RNA-seq levels (2078 vs. 2295 rpkm in WT vs. Δ*pnp*) but very different Φ_end_ values (0.48 vs. 0.10) (Fig. 5A-B). Globally, the log fold-change in RNA-seq signal is closely centered around 0, but Φ_end_ values decrease by 3-fold on average. This result indicates that RNA-seq fails to report the substantially decreased levels of functional mRNA, which are accompanied with increased partial transcripts that are indistinguishable by RNA-seq.

**Figure 5:**
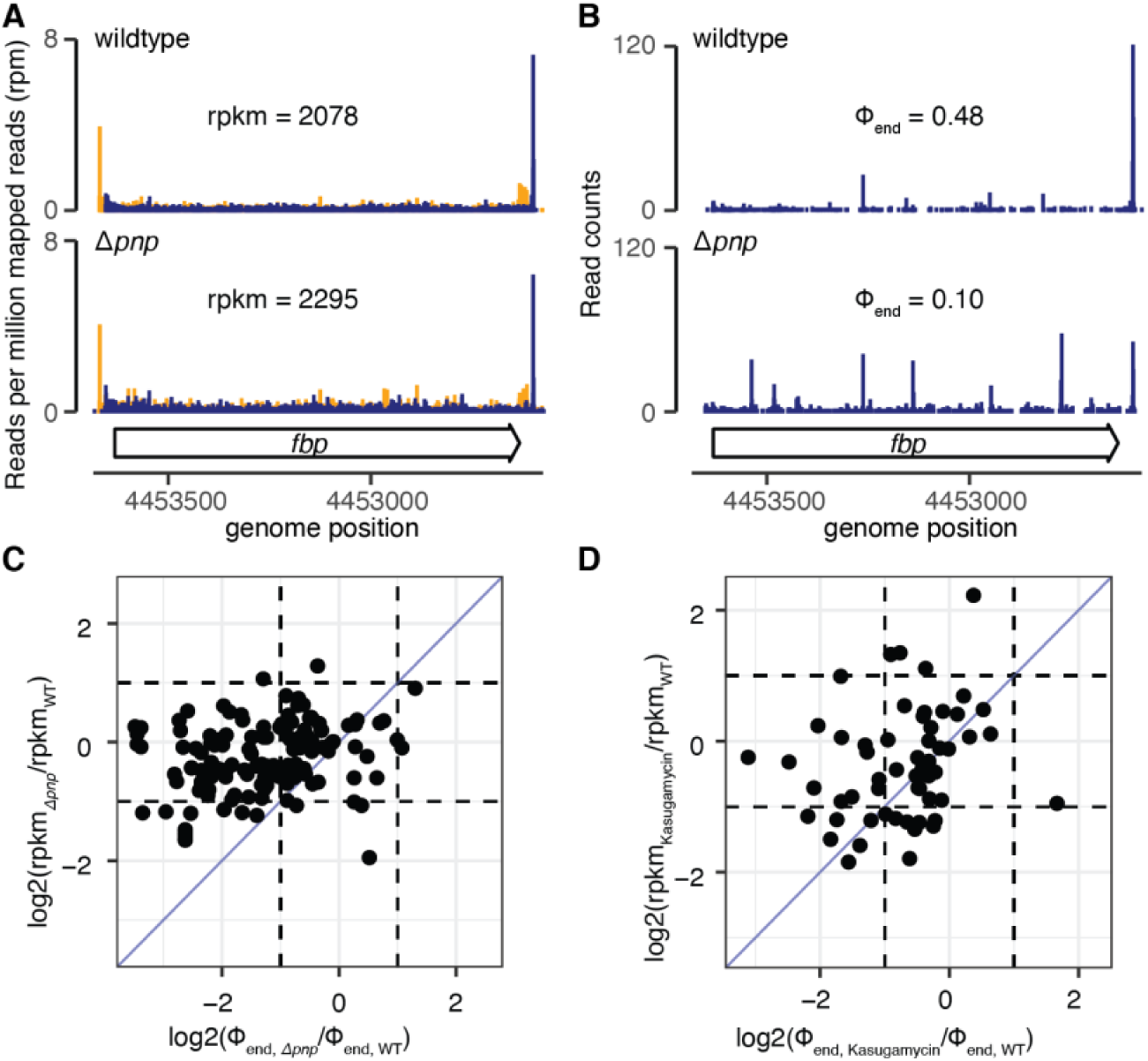
Transcriptome heterogeneity confounds quantification of functional mRNAs by RNA-seq. (A) Representative RNA-seq data in wildtype *E. coli* and Δ*pnp*. Rend-seq data show coverage across *fbp* (colored as in Fig. 1D). The reads per kilobase mapped per million reads (rpkm) in each sample are calculated using 3′-mapped reads, excluding the first and last 50 nts of the gene. (B) Representative 3′ end sequencing data for the same samples as in (A). Φ_end_ values are indicated. (C-D) Fold-changes in Φ_end_ and rpkm values for simple TUs between Δ*pnp* and wildtype (C) and sample with and without Kasugamycin treatment (D). Each dot represents one transcription unit. Dashed lines correspond to 2-fold changes in either Φ_end_ or rpkm, whereas blue lines mark the diagonal that would indicate the same degree of change by both metrics. In (D), cells were treated with the translation inhibitor Kasugamycin for 15 min.

Consistent with the global increase in partial, non-functional transcripts in the Δ*pnp* strain, we found that tmRNA, which rescues stalled ribosomes at the end of RNAs that lack stop codons (Keiler, 2015; Müller et al., 2021), has increased association with ribosomes (Fig. S5B-E). Meanwhile, the cellular backup system mediated by the alternative ribosome rescue factor ArfA (Chadani et al., 2011, 2010) is strongly upregulated (Fig. S5F), suggesting a pronounced need for ribosome rescue due to accumulation of decaying transcripts. These results, together with a much longer cell doubling time (Fig. S5A), corroborate the substantially different functional transcriptome that was not captured by RNA-seq.

Similar to cells lacking PNPase, we found that other genetic or antibiotic perturbations also lead to changes in the functional transcriptome without co-occurring expression changes measured by RNA-seq (Fig. 5D, S5G-I). In other words, RNA-seq largely underestimated the pronounced effect on RNA integrity that is visible in Φ_end_ changes. In conclusion, our study shows that end-based mapping approaches are important to study RNA decay and that prevalent decay intermediates can interfere with gene expression profiling.

## Discussion

We devised an approach to obtain a molecular representation of *in vivo* transcriptome that reflects their life cycles and capabilities for translation. We find that in *E. coli*, mature full-length mRNAs are often not the dominant species. Most partial transcripts are *in vivo* RNA decay intermediates, albeit they are less prevalent in the ribosome-bound transcriptome. We showed that the presence of decay intermediates in total RNA can distort quantifications for the abundance of full-length, functional mRNAs.

By profiling individual 3′ ends of cellular RNA, we obtain a single-nucleotide perspective of transient RNA decay dynamics that are possibly linked to other aspects of gene regulation. 3′ ends generated by endonucleoytic RNase cleavage are unstable in *E. coli* due to the presence of 3′-5′ exonucleases (Cohen and McDowall, 1997; Hui et al., 2014; Mohanty and Kushner, 2016; Bechhofer and Deutscher, 2019; Trinquier et al., 2020). Hence, we were initially surprised to pick up such cleavage signatures, but it highlights that our approach captures snapshots of *in vivo* dynamics and transient intermediates of RNA decay. Consistent with previous studies, we detected a sizeable fraction of adenylated transcripts (Maes et al., 2017; Mohanty and Kushner, 2006) and found that very short tails of non-templated As are present at RNA 3′ ends at steady-state (Waters et al., 2017). The length of A-tails likely reflects the balance of concomitant adenylation and deadenylation by poly(A) polymerase PcnB and exonucleases, respectively.

Changes to transcriptome heterogeneity are conceivable in conditions where mRNA decay is perturbed, which would in turn lead to translational stress. Upon deletion of non-essential enzymes involved in mRNA decay in *E. coli* or inhibition of translation initiation (Schneider et al., 1978), we detected decreases in Φ_end_, with the strongest effect observed for the deletion of PNPase, which also showed a strong growth defect (Fig. 5, S5). We found that the increases in partial transcripts upon PNPase deletion are accompanied with accumulation of stalled ribosome on non-stop RNAs, as indicated by the increased demand for ribosome rescue by *trans*-translation and through the alternative ribosome rescue factor *arfA* (Fig. S5). Similar responses have been detected when depleting the ribosome recycling factor RRF or overexpression of the ribosome-associated mRNA cleaving toxin RelE (Hwang and Buskirk, 2017; Saito et al., 2020). In general, environmental responses and stress conditions might result in changes of transcriptome heterogeneity, e.g., due to altered mRNA decay (Vargas-Blanco and Shell, 2020), that manifest themselves in a pronounced ribosome rescue response.

The comparison between RNA-seq and 3′ end sequencing data upon PNPase deletion reveal limitations in RNA-seq to quantify functional mRNAs (Fig. 5). Conventional RNA-seq does not distinguish between partial and full-length transcripts and thus reports average expression values that can arise from multiple possible combinations of partial and full-length transcripts. Many partial transcripts are longer than the RNA size cutoffs that are commonly used during purification, such as the 200-nt cutoff in many column-based RNA extraction methods (Fig. 1E, S1, 3C). Thus, this population can substantially contribute to the expression estimate despite being non-functional for protein production in many cases. Similar limitations may apply to other RNA quantification methods, such as RT-qPCR which is based on amplification of ∼100-nt regions. Future improvements to full-length RNA sequencing approaches, especially mitigations to current length biases, will greatly advance our ability to quantify the functional part of the transcriptome.

The high transcriptome diversity due to RNA decay intermediates requires consideration in all biological systems. In bacteria specifically, the life of an mRNA is shaped by co-occurrence of all steps of gene expression with little spatial separation in the cytoplasm, and thus with ample opportunities for crosstalk and interference. We quantified the resulting transcriptome distribution in this study in the two bacterial model systems *E. coli* and *B. subtilis*. Similarly in eukaryotes, transcriptome diversity due to RNA decay intermediates has been previously identified and utilized to reveal translation dynamics (Pelechano et al., 2015), but the molecular census remains to be quantified exactly. Furthermore, in mitochondria and plastids, gene expression is subject to similar constraints as in bacteria. Hence, the observed heterogeneity of cellular mRNAs may play important roles in gene expression and cell physiology across species.

## Materials and methods

### Strains, cell growth and harvest

Strains used in this study are listed in Table S1. To generate fresh gene deletions in the strain background MG1655, P1 transduction from the KEIO collection was done as described previously (Baba et al., 2006; Thomason et al., 2007). To harvest cells in exponential growth, cells were backdiluted from overnight cultures into the respective growth media to enable ∼10 doublings at 37°C prior harvest at an OD_600_ smaller or equal to 0.3 (Sezonov et al., 2007). *E. coli* cells were grown in LB or MOPS Complete media supplemented with 0.2% Glucose. *B. subtilis* cells were grown in MCCG media (Parker et al., 2020). For total RNA analysis, 5 ml of cells were transferred into 0.55 ml of ice-cold stop solution (95:5 Ethanol:Phenol) in a 15 ml conical tube, quickly inverted and spun at 3220 g at 4°C for 5-10 min. The supernatant was discarded and cell pellets flash-frozen in liquid nitrogen and stored at −80°C until lysis and RNA extraction. For nascent RNA and polysome analysis, 250 ml of cells were filter harvested at 37°C and scraped into liquid nitrogen (Johnson and Li, 2018). The cell paste was kept at −80°C until lysis and RNA extraction.

### Genome version, annotation and simple transcription unit definition

For all experiments described here, the NCBI Reference Sequence and CDS annotation for the *E. coli* genome NC_000913.2 and the *B. subtilis* genome NC_000964.3 were used.

#### Definition of simple transcription units

We focused our analysis and quantification of transcriptional and decay intermediates to simple transcription units (TUs). Simple TUs are transcripts with only one isoform – i.e. have only one transcription start site (giving rise to the mature 5′ end) and one transcription termination or processing site (giving rise to the mature 3′ end). In order to classify regions in the *E. coli* and *B. subtilis* genome as simple TUs we utilized high-coverage Rend-seq data from *E. coli* MG1655 grown in the same conditions as for our 3′ end-seq experiments (MOPS-Complete and LB, harvested at OD_600_ 0.3) and *B. subtilis* 168 grown in LB until OD_600_ 0.3 (Lalanne et al., 2018)(GSM2500131, GSM2500127, GSE189181). By enriching for mature 5′ and 3′ ends of RNA, Rend-seq allows to map transcript isoform ends (peaks) with single nucleotide precision (Lalanne et al., 2018). We called 5′ and 3′ peaks as described previously (Lalanne et al., 2018), using a threshold Z-score of 12. We considered 5′ or 3′ peaks that were within +/- 2 nt in different datasets as 1 peak, because this corresponds to the expected peak width from these data and can have maxima that vary by +/- 2 nt around the center (Lalanne et al., 2018).

We then identified all transcripts which met our criteria for being a simple TU. First, we matched any 5′ and 3′ peaks that were connected by expressed regions and separated by regions that did not have sufficient read coverage (defined by median coverage). This procedure creates a list of potential mRNA isoforms across the genome. Second, we added the restriction that the size of each transcription unit lies within the 2 - 98% quantile of all isoforms (75 - 20872 nt for *E. coli*). This resulted in 7635 isoforms for *E. coli* from 3617 unique 5′ peaks and 1681 unique 3′ peaks. Third, we excluded rRNA and tRNA regions from further analysis by keeping only non-rRNA and tRNA-overlapping isoforms using bedtools intersect (rRNA & tRNA annotation Table S3, bedtools/2.25.0 (Quinlan and Hall, 2010)). That reduced the number of isoforms to 5676. Fourth, to obtain a candidate set for simple TUs, we overlapped the isoforms with each other using bedtools intersect and only kept the 215 isoforms that do not overlap with any other ones. Lastly, we manually inspected and curated the simple TUs in the genome browser IGV and MochiView to ensure that no apparent peaks in Rend-seq data (that did not meet our stringent peak calling criteria) were in the vicinity. These would indicate less abundant transcript isoforms (Homann and Johnson, 2010; Robinson et al., 2011). This reduced the number of simple TUs to 142. These simple TUs were used for the 3′ end analysis reported in the main text (Table S4).

The same approach (omitting the curation in the genome browser) yielded 418 simple TUs in *B. subtilis* that were used for the analysis shown in Fig. S3 (Table S5). It is likely that there are more simple TUs that we did not capture using our approach due to our conserved criteria for RNAs collected in exponential phase in rich media conditions. Furthermore, many 3′ ends of transcription units in *E. coli* are less well-defined compared to 5′ ends, and thus some 5′ ends cannot be confidently linked to a 3′ end.

#### Definition of genomic regions encoding mRNAs

For analysis of 3′ and 5′ ends profiles with regard to RNase E cleavages, RNA polymerase pauses, and non-templated adenylation in any protein-coding gene, we developed a coarse definition of genomic regions that encode mRNAs. To this end, we grouped CDSs based on their distance to each other (<121 nt) and strand orientation. To include untranslated regions (UTRs) of mRNAs, we extended CDSs in each direction by 60 nt, which is the typical UTR length observed in Rend-seq (Table S6). These genomic regions can have multiple mRNA isoforms within.

### Synthesis of reference RNAs for spike-in

We used a total of five reference RNAs to spike-in to our 3′ end sequencing samples for quality control purposes. The two shortest reference RNA (28 nt and 60 nt) were ordered as PAGE-purified RNA oligos from IDT (Table S2). The remaining three RNA spike-ins were generated by *in vitro* transcription (details below).

We generated the 120-nt reference RNA by amplifying the *S. cerevisiae* gene PDR5 from genomic DNA using oligos oLH245 and oLH238 (Table S2). We followed the PCR with an agarose gel extraction and a column purification using the Oligo Clean & Concentrator kit (Zymo Research). The DNA template was then *in vitro* transcribed using the HiScribe T7 high yield RNA synthesis kit (NEB) in a 20 μl reaction for 10 h at 37°C. We then digested the template DNA using TurboDNase at a concentration of 0.1 U/μl in 200 μl and cleaned up the reaction using the RNA Clean & Concentrator-25 kit (Zymo Research). Finally, we performed a gel extraction from a 10% polyacrylamide gel as described in (Johnson and Li, 2018) and quantified the resultant RNA concentration using the fluorometric Qubit RNA BR assay.

To generate the 920-nt reference RNA, the pTXB1vector (NEB) was digested with BamHI, gel purified and *in vitro* transcribed using the HiScribe T7 high yield RNA synthesis kit (NEB) in a 20 μl reaction for 10 h at 37°C. TurboDNase digestion with a final concentration of 0.1 U/μl in 200 μl, clean-up using the RNA Clean & Concentrator-25 kit (Zymo Research), agarose gel analysis and quantification with the Qubit RNA BR assay followed.

To obtain a 1800-nt reference RNA, the FLuc Control Template from the HiScribe T7 high yield RNA synthesis kit (NEB) was *in vitro* transcribed and purified as described the linearized pTXB1 vector.

Aliquots of each reference RNA were kept in 10 mM Tris pH 7.0 at the final concentration for pooling at −80°C (1 reference RNA set for 1 RNA extraction contained approximately 6 ng 28-nt reference RNA, 57 ng 60-nt reference RNA, 53 ng 120-nt reference RNA, 70 ng 920-nt reference RNA, 170 ng 1800-nt reference RNA). Each pooled batch was sequenced by 3′ end-sequencing without spiking it into a cell suspension to serve as an input for quality normalization (4 input replicates in Fig. 1C, normalized to mean Φ_end_ of these 4 technical replicates).

### RNA purification

For all total RNA extractions from cell pellets, cell pellets were thawed for 4 min in a 4°C pre-cooled tabletop centrifuge while spinning at 3220 g. Tubes were transferred on ice and residual media was removed by pipetting.

#### Acid-Phenol:Chloroform extraction (Hot phenol)

500 μl Acid-Phenol:Chloroform, pH 4.5 (with IAA, 125:24:1, Thermo Fisher Scientific) containing 29 μl of 20% SDS was prewarmed to 65°C prior adding it to 500 μl sample. The sample solution containing Acid-Phenol:Chloroform and SDS was incubated for 5 min at 65°C in an Eppendorf Thermomixer at 1400 rpm. Subsequently the solution was transferred to ice and chilled for 5 min. After a 2 min spin at room temperature and 20,000 g 90% of the aqueous layer were transferred to a fresh tube and precipitated with Sodium Acetate / Isopropanol. The sample was further purified using the RNA Clean & Concentrator-5 kit (Zymo Research) and DNase treated subsequently (details in ‘RNA cleanup’ section).

#### RNAsnap

RNA extraction using RNAsnap was done as previously described (Stead et al., 2012) with small adjustments. The cell pellets were resuspended in 500 μl RNA extraction solution (RES, 18mM EDTA, 0.025% SDS, 1% BME, 95% Formamide, RNA spike-ins). The RES was prepared fresh each time from stock components and reference RNA aliquots were added just before use. The RES-cell suspension was vortexed for 50 s and incubated for 7 min at 95°C. For *B. subtilis* cells, the RES-cell suspension was added to 0.2 ml cold 0.1 mm Zirconia beads, vortexed for 5 min at maximum speed and incubated for 7 min at 95°C. A 5 min spin at room temperature at 20,000 g followed. 400 μl of supernatant were transferred to a fresh tube containing 1.6 ml of DEPC-water. Subsequently, this 1:4 dilution was split into 4x 500 μl aliquots for sodium acetate / isopropanol precipitation (details in ‘RNA cleanup’ section). Right after precipitation the sample was further purified using the RNA Clean & Concentrator-5 kit (Zymo Research) and DNase treated subsequently.

#### RiboPure RNA Purification Kit

RNA extraction with the RiboPure RNA Purification Kit (Thermo Fisher Scientific) was done as recommended by the manufacturer.

#### RNeasy RNA Purification Kit

Cell pellets were resuspended in 100 μl of 4 mg/ml lysozyme in 10 mM Tris, pH 8.0 with reference RNA and incubated for 5 min at 37°C. RNA was extracted as described in the manual without the use of gDNA columns. Instead, the sample was DNase treated and purified as described below. rRNA was removed from 20 μg of RNA with 2 reactions of the MICROB*Express* Bacterial mRNA Enrichment Kit (Thermo Fisher Scientific).

#### Trizol extraction

Cell pellets were resuspended in 100 μl of 4 mg/ml lysozyme in 10 mM Tris, pH 8.0 with reference RNA and incubated for 5 min at 37°C. 1.5 ml Trizol was added and mixed by inversion. After 5 min incubation at room temperature, 300 ul of chloroform were added. After vigorous shaking for 15-30 s the lysate was spun for 15 min at 12,000 g in a pre-cooled 4°C centrifuge. About 800 μl of the aqueous phase were transferred to a fresh tube and precipitated with sodium acetate / isopropanol. As for RNAsnap, the sample was further purified using the RNA Clean & Concentrator-5 kit (Zymo Research) and DNase treated subsequently.

### RNA cleanup by precipitation and column and DNase treatment

To concentrate and cleanup RNA samples, Sodium Acetate - Ethanol or Isopropanol precipitation was carried out. 3 M Sodium Acetate (pH 5.4, Life Technologies) was added to make up 1/10th of the sample volume. 1-2 μl of GlycoBlue (Invitrogen) was used as co-precipitant. Depending on the sample volume either 3 volumes of ice-cold 100% Ethanol (<300 ul) or 1 volume of ice-cold 100% Isopropanol was added. After thorough mixing, samples were precipitated at −80°C for at least 30 min and spun for at least 60 min in the pre-cooled 4°C centrifuge at 18213 g. RNA pellets were washed with at least 250 μl of ice-cold 80% Ethanol and spun for at least 4 min. After aspiration by pipetting RNA pellets dried at room temperature for 3-5 min and were resuspended in 10 mM Tris, pH 7.0. To clean up samples further and remove traces of short DNA oligos post DNase treatment, we purified RNA samples by RNA Clean & Concentrator-5 kit (Zymo Research). If this kit was used prior DNase treatment, elution was done with 85 μl DEPC-water. Otherwise, samples were eluted into 10 mM Tris, pH 7.0. For DNase treatment, 5 μl 10x Turbo DNase buffer and 5 μl 2U/μl TurboDNase (Thermo Fisher Scientific) were added to 85 μl of sample and incubated for 20-30 min at 37°C. RNA was cleaned up either with the RNA Clean & Concentrator-5 kit (Zymo Research) or by Ethanol precipitation.

### Protein analysis

Protein samples were run on a 4-12% SDS polyacrylamide gel followed by SyproRuby staining (Thermo Fisher Scientific) and Western Blotting (as described in the NuPAGE Technical Guide from Thermo Fisher Scientific). Western blots of different fractionations during the RNA polymerase immunoprecipitation and polysome gradient fractionation were performed to assay efficiency of flag-tagged RNA polymerase enrichment using the monoclonal Anti-Flag M2 antibody from mouse (F1804, Sigma).

### RNA polymerase immunoprecipitation

The RNA polymerase immunoprecipitation was carried out as described in (Larson et al., 2014) with the following modification. Nascent RNAs were extracted from the magnetic-bead bound fraction by adding 1.5 ml Trizol to the sample and proceeding as described above. For comparison to total RNA, 100 μl of lysate were diluted to 250 μl in lysis buffer and RNA was extracted using Trizol. Following DNase treatment and sample cleanup, rRNA was removed using the MICROB*Express* Bacterial mRNA Enrichment Kit (Thermo Fisher Scientific) and precipitated with isopropanol. RNA levels were quantified fluorometrically with the Qubit RNA BR assay. 1.8 μg of RNA was used for 3′ end-seq library preparation and 3-5 μg for Rend-seq library preparation.

### Polysome gradient centrifugation

Cell paste from 250 ml of cells filter-harvested at or below OD_600_ 0.3 was cryo-lysed as done for ribosome profiling (Johnson and Li, 2018). 1 M sodium chloride (Mohammad et al., 2019), 200 U/ml SUPERaseIn RNase Inhibitor and 100 U/ml TurboDNase were included in the lysis buffer in addition to the standard composition of 20 mM Tris pH 8.0, 100 mM ammonium chloride, 0.4% Triton X-100, 0.1% NP-40, 10 mM magnesium chloride and 5 mM calcium chloride. 250 μl of lysate were loaded onto a 10-55% sucrose gradient and spun for 156 min at 151263 g with a SW41 Ti rotor at 4°C. 12 gradient fractions were collected into disposable glass tubes for fraction collection with the same instrumentation as for ribosome profiling (Johnson and Li, 2018). The remaining liquid and pellet in the tube were either taken as a separate fraction (Fig. S5C) or pooled as a final fraction (Fig. S5B). Each fraction was transferred to a fresh tube and precipitated with ice-cold isopropanol as described above. The pellet was resuspended in 500 μl 10 mM Tris, pH 7.0 containing 0.5% SDS and RNA was extracted subsequently using hot phenol extraction as described above. Following DNase treatment and sample cleanup, RNA levels were quantified fluorometrically with the Qubit RNA BR assay and 3-5 μg were used for Northern blot analysis and 1.8 μg for 3′ end-seq library preparation.

### PCR & RT-qPCR

Reverse transcription (RT) of RNA was done using SuperScript III reverse transcriptase (Thermo Fisher Scientific). The protocol recommended for the enzyme was used in either 10 or 20 μl reactions. 100 ng of RNA were used as input. Random hexamers (Thermo Fisher Scientific, 1 μl per 20 μl) were used for priming. A ‘no RT primer’ and a ‘no enzyme’ control were also included to assess random priming from residual short gDNA oligonucleotides and presence of contaminating gDNA, respectively. RNA, dNTPs and RT primer were denatured at 80°C for 3 min and immediately cooled on ice for at least 1 min. Afterwards RT buffer, 0.1 M DTT and SuperaseIn (Thermo Fisher Scientific) were added in the respective amounts. Superscript III enzyme was added last (or water for ‘no enzyme’ controls). Samples were incubated at room temperature for 5 min, then 55°C for 45 min. RNA was hydrolyzed at 95°C with 0.1 M NaOH (final concentration) for 5 min and neutralized with HCl. cDNA samples were diluted 1:10 for further use in qPCR. For subsequent qPCRs, Kappa qPCR reaction mix (Roche) was used. In each well of a 96-well plate we added 5 μl 2x Kappa qPCR reaction mix, 3 μl of the 1 μM primer solution and 2 μl of the 1:10 diluted cDNA solution.

We assayed each sample in combination with several controls, such as a ‘no RT primer’, ‘no enzyme’ and a ‘no cDNA’ control, in 2-3 technical replicates. Each run was done as recommended by the manufacturer. Only samples, which had a Ct value at least 10 cycles lower than the control Ct value were used for further analysis. Especially, a smaller than 10 Ct value difference to the ‘no enzyme’ control suggests residual genomic DNA that needs to be reduced by a second round of DNase digestion prior RT. Fortunately, TurboDNase treatment as described above usually removes most genomic DNA and small fragments of DNA that could prime as random primers (detectable by ‘no RT primer’ control) are cleared away efficiently by the RNA Clean & Concentrator-5 kit (Zymo Research). Hence, a second round of DNase treatment was never necessary in our hands.

To quantify gene expression differences the double-Ct method was employed (Rao et al., 2013). Namely, Ct values were first normalized by the expression of *cysG*, a gene whose expression level does not change across the conditions assayed and that was used in previous analyses (Pobre and Arraiano, 2015). After normalizing both the target strain and the wildtype strain to *cysG* we then compared the ratios of their relative abundance to estimate fold changes in the gene of interest between strains.

For conventional PCRs, Q5 DNA polymerase (NEB) was used with recommended reaction settings and reagent concentrations. 35-40 cycle PCR reactions were carried out. PCR products were analyzed by conventional 1.5% Agarose gel electrophoresis. Templates for *in vitro* transcription reactions were gel purified using the QIAquick Gel Extraction kit (Qiagen).

### Northern blot

Northern blot analysis was done as described in (Lalanne et al., 2018) and DNA oligos used as probes are given in Table S2. For total RNA analysis 2% agarose gels were used. 1 pmol of DNA probe was used for 5′ terminal γ-^32^P labeling followed by cleanup with ProbeQuant G-50 columns (Sigma) when assessing mRNA levels. To probe highly abundant RNAs, such as *ssrA* and *rnpB* 0.01 pmol of DNA probe sufficed and enabled full removal of signal by stripping of transcript specific probes with three 20 min washes with boiling 0.1% SDS followed by an overnight incubation at 47°C.

Labeled membranes were exposed to a phosphor storage screen (GE Life Science) for hours to days depending on the signal strength and imaged with a 635 nm laser scanner (Typhoon FLA9500, GE Life Sciences or Amersham Typhoon) with 750 V and 100 μm pixel size. By testing several exposure times of the radioactively labeled membranes to the phosphor storage screen and testing different PMT voltages in the range from 400-750 V we ensured that neither the phosphor storage screen nor the Typhoon scanner saturated (none of our tested conditions yielded saturation). The Amersham Typhoon Scanner Control Software 2.0.0.6 converts the original 32-bit data into a 16-bit data format for saving as tif-file that was used for further analysis in Fiji (Schindelin et al., 2012). Images were cropped to only include the membrane of interest and intensity profiles plotted per lane with a linewidth of 50 pixels. These images are shown in Fig. 1E, S1A, 3C, S5D. For all northern blot images, we show the full range of pixel intensities. The fraction of full-length RNAs was quantified semi-automatically for Northern blots that showed uniform background signal across the full membrane and showed uniform sample and run characteristics for all lanes on the gel. The median background signal above the full-length signal was subtracted. The Northern blot shown in Fig. 1E showed a background distribution that increased linearly from top to bottom. Hence, we determined the median background signal above the full-length product and at the bottom of the blot, where no RNA was present and subtracted the linearly increasing profile between these two background intensities.

Upon log-transformation of the background-subtracted intensities for all lanes on one membrane, 2 inflection points, indicating the boundaries of full-length RNA to 1) no signal above and 2) partial transcript signal below, were defined. The signal between these two points reflects full-length RNA signal (FL) (shading in Fig. 1F). The signal below the 2^nd^ inflection point reflects any signal of partial transcripts (P). The ratio of FL / (FL+P) was defined as the fraction of full-length RNAs F_FL_ and compared to Φ_end_ derived from 3′ end-seq (Fig. 1G).

For the analysis of *ssrA* distribution across polysomes (Fig. S5B-D), Northern blots were performed for three biological wildtype replicates (representative wildtype replicate shown in Fig. S5B) and one replicate for *Δpnp* (Fig. S5C). To quantify the increased signal in polysomal fractions of *Δpnp* that is already visible by eye, we calculated the background subtracted cumulative signal per equally sized band and adjusted for the amount (μg) of loaded RNA per lane. Individual fractions between gradients are often slightly differently aligned with respect to the migration of individual ribosomal subunits, monosomes and polysomes. Hence, we did not compare individual fractions, but grouped fractions as ribosome-free (subunits & lightest fraction(s); fraction 1,2 and 0-2, respectively in WT and *Δpnp*) and ribosome-associated fractions (fractions >2). We summed the background- and loading-adjusted band intensities for both groups and show the relative proportions in Fig. S5D.

### RNA-seq and Rend-seq

Oligos used for library preparations are listed in Table S2. RNA-seq for expression quantification in Fig. S5G-I was executed as described in (Parker et al., 2020). Rend-seq libraries were prepared as described in (Lalanne et al., 2018) from 20 μg of RNA prior rRNA removal. RNA fragmentation was done for 25 s. Final libraries were analyzed for size, quality and concentration using qPCR and the Fragment analyzer at the Biomicrocenter at MIT and sequenced on Illumina HiSeq 2000 or NextSeq500 machines. RNA-seq and Rend-seq data processing and mapping was done as described in (Lalanne et al., 2018; Parker et al., 2020).

### 3′ end-sequencing library preparation

For 3′ end sequencing, the Rend-seq protocol was modified in the following ways: 3′ end ligation with Linker-I was done with 1.8 μg of non-fragmented total or nascent RNA, followed by RNA precipitation using isopropanol. This ligation targets hydroxylated RNA 3′ ends that are typical products of transcription and canonical RNA decay (Yang, 2011). Phosphorylated RNA 3′ ends that may arise as a result of non-enzymatic fragmentation or RNase A contamination are not ligated using this approach (Herzel, 2015; Yang, 2011). RNA was fragmented for 2 min following ligation, which was followed again by RNA precipitation using isopropanol. 32-63 nt fragmented 3′ end-ligated RNA was size selected from a 15% TBE-Urea polyacrylamide gel and precipitated for reverse transcription, circularization and final PCR amplification with 6-8 cycles as described in (Lalanne et al., 2018).

Our approach differed from Term-seq (Dar et al., 2016) in the following ways: RNA ligation was done with truncated T4 RNA ligase II, K227Q instead of T4 RNA ligase 1 and different adaptor sequences. For cleanup between reaction steps RNA and cDNA was precipitated or gel extracted & precipitated, whereas sample cleanup in Term-seq was done with SPRI beads. Primers and enzymes for reverse transcription and consequently buffers and reaction conditions were also different. To add PCR handles on both sides of the cDNA fragment, our protocol included an intramolecular cDNA circularization step and Term-seq relied on an intermolecular ligation of a separate DNA adaptor. By design, our samples were sequenced from the 5′ end of the insert into the 3′ end adaptor and Term-seq sequences from the 3′ end.

### 3′ end-seq processing and mapping

In analogy to Rend-seq, 3′ end linker sequences were trimmed. To deal with non-template addition during reverse transcription, all reads were further trimmed with the fastx_trimmer of the fastxtoolkit/0.0.13 by 1 nt at their 5′end and only reads of 15 nt length or longer were kept for mapping with bowtie/1.2 using the options -v 0 -k 1 (Langmead et al., 2009). For further data analysis mapped data were transformed into bam-format and bedgraph-format using samtools/1.5 and bedtools/2.25.0 (Li et al., 2009; Quinlan and Hall, 2010). Coverage data include only 3′ terminal single nucleotide counts per read. To identify 3′ adenylated transcript ends, the unmapped reads were trimmed using cutadapt/1.16 (Martin, 2011). First, 3′ terminal CCAs were removed with the following settings: -a “CCA$” --minimum-length 15 -O 3 --trimmed-only. This step is necessary to avoid confusing CCA-tails, e.g. at tRNA 3′ ends with 3′ end adenylation that arise through different mechanisms *in vivo* (Wellner et al., 2018). Second, A-tails of any length were trimmed (-a “A{38}” --minimum-length 15 -O 1 --trimmed-only). These trimmed reads were mapped and further converted with the same settings and code as untrimmed reads above. For total RNA 3′ end analysis mapped trimmed (i.e. A-tailed) and untrimmed 3′ end coverage profiles were summed.

### Reference RNA mapping

Unmapped reads after mapping 3′ end-sequencing to the *E. coli* genome (described above) were mapped with the same settings and code as above, but to the five reference RNA template sequences (Table S2). In most cases T7 RNA polymerase adds a few non-templated nucleotides at the 3′ end of the template (Kao et al., 1999). To include these 3′ ends for full-length RNA analysis, unmapped reads after mapping to the reference RNA template were five times iteratively trimmed by 1 nt, filtered for length to only include reads of 15 nt length or longer (fastx_trimmer -Q 33 -t 1 -m 15) and remapped to the template. After trimming 5 nt only background mapping was observed and thus trimming was stopped. Trimmed mapped reads and reads that mapped to the reference RNA template prior trimming were merged and converted to bedgraph files using sam- and bedtools.

### Expression, Φ_end_ calculation, rRNA ratio calculation

To quantify mRNA levels (rpkm, reads per kilobase million) per coding sequence (CDS) from Rend-seq data, coverage at each base was divided by the total number of mapped reads in million. This yielded reads per million (rpm). Next, we calculated the total coverage per CDS excluding 50 nt at both ends of the CDS to avoid conflating effects by end-enrichment in Rend-seq and divided by the length of the CDS in kb minus 0.1 kb, yielding rpkm. Depending on the depth of the RNA-seq dataset we either required a minimum of 20 reads or 100 reads to consider a rpkm for further analysis.

The fraction of mature 3′ ends Φ_end_ was calculated as the ratio of 3′ ends mapping to the mature 3′ end of simple TUs (+/- 2 nt based on the expected possible peak width (Lalanne et al., 2018)) and all 3′ end counts mapping to that TU (Table S7).

The three very differently sized ribosomal RNAs (rRNAs) 5S (0.12kb), 16S (1.5kb) and 23S (2.9kb) are expressed at a molar stoichiometry of 1 and very stable with half-lives beyond the doubling time of *E. coli* (Bremer and Dennis, 1996; Dong et al., 1996). Hence, we reasoned that these can be used as internal size standards and their mature 3′ end signal should reflect the 1:1 ratio in samples that were not depleted of rRNAs prior library preparation. That is the case for our RNAsnap and hot phenol extracted RNA samples (Fig. 1B). In addition to this analysis, Φ_end_ of rRNAs are also high above 0.8 even for the 23S rRNA (Fig. 3A).

### Peak calling

In 5′ end and 3′ end sequencing datasets, 3′ end or 5′ end coverage peaks are called based on Z-score calculation across a window of 100 nt. We used peak calling in the analyses to identify RNA polymerase pause sites (Fig. 2), putative RNase E sites (Fig. 3) and mature 3′ ends across all TUs (Fig. 3F). Specifically, we extracted the read coverage in a region +/- 50 nt of any position covered by a read in the dataset and normalized by the total amount of reads in this window. We required a minimum of 15 reads in each window for Z-score calculation. To calculate mean and standard deviation of the background the center position was excluded. The Z-score reflects the normalized coverage at the position of interest minus the background mean divided by the standard deviation. Z-scores above 7 were considered peaks. These could be within or outside of CDSs.

To call putative RNase E sites from remapped 5′ end sequencing data from (Clarke et al., 2014), we compared data in the rne-3071 ts strain (temperature-sensitive mutant of RNase E) and the wildtype strain. 5′ end peaks should be absent or substantially reduced in the rne-3071 ts strain compared to wildtype. Hence, we required that Z-scores in the wildtype strain were above 7 and at least four times lower in the 5′ end-seq data in the rne-3071 ts strain. Lastly, we focused our analyses on sites within protein-coding transcription units. The RNase-dependent sites are listed in Table S9.

### Quantification of nascent RNA fraction in total RNA

For the nascent RNA 3′ end data, 21355 positions had Z-scores above the cutoff of 7 and sufficient read coverage that were 2.1% of all positions passing read cutoff of 15 in the window around the position of interest. We overlayed peaks with the CDS annotation using bedtools intersect and classified them according to their relative position as in- or outside of CDSs (71% inside CDSs). To visualize the nucleotide distribution around peaks internal to CDSs, the underlying sequence 10 nt upstream and 5 nt downstream around peak positions was extracted from the genome fasta sequence and visualized as sequence logo with Weblogo 3 (Crooks et al., 2004).

To refine the classification of RNA polymerase elemental pause positions for the quantification of nascent RNAs in total RNA, we calculated a pause motif score across the 16 nt of interest and exclude the 25% lowest scoring sites from any further analysis. To this end, we calculated the position weight matrix from all sites in CDSs that have a peak with a Z-score greater 7. From that, we generated a log-odds matrix by comparing to the nucleotide frequencies within protein-coding genes. For each individual 16 nt peak sequence, the pause motif score was calculated by summing up log-odds matrix entries at the respective sequence position (column) and base identity (row). Peaks with pause scores below 2 (25% lowest scoring peaks) did not show the sequence preference as given in Fig. 2C when visualized as sequence logo and were not used for the quantification of nascent RNAs in total RNA that is described below.

The peak distribution and sequence comparison between total and nascent RNA sample suggested little nascent RNA in the pool of total RNA (Fig. 2B-C). To estimate the nascent RNA fraction in total RNA, we reasoned that the amount of total RNA 3′ ends at positions of RNA polymerase elemental pause sites provides an estimate of the contribution of nascent RNAs to the total transcriptome. However, not all nascent RNA 3′ ends fall into these pause sites. Hence, we calculated the fraction of nascent RNA signal in pause sites f_pause_ compared to the whole transcription unit (total TU counts); the same was done for total RNA (without the adjustment for biochemical enrichment, see below). The ratio of the two fractions (total/nascent) was considered to reflect the “Nascent RNA fraction in total RNA”, which is an upper bound for nascent RNA in the pool of total RNAs, because also 3′ ends at pause positions could come from decaying transcripts that happen to be caught at that position.

Some 3′ end signal across the whole transcription unit originates from total RNA due to the incomplete biochemical enrichment of nascent RNAs. To account for this, we multiplied the total TU counts with 0.91, which is the median of the internal end fraction for simple TUs with 30 or more 3′ end reads in the nascent RNA sample. It indicated that at least 9% of 3′ ends came from total RNA. The distribution of the “Nascent RNA fraction in total RNA” is given in Fig. 2D and Table S8 for simple TUs and all TUs (Table S6, mRNA encoding regions). The medians of these distributions are 0.14 and 0.16, respectively.

### Analysis around putative RNase E sites, 3′ end peaks and across CDSs

#### Sequence logo of RNase E sites (Fig. 3B)

We aimed to visualize the nucleotide distribution around putative RNase E sites. To this end, we extracted the underlying sequence 5 nt upstream and 3 nt downstream at peak positions from the genome fasta sequence using bedtools getfasta and visualized them as sequence logo with Weblogo 3 (Crooks et al., 2004; Quinlan and Hall, 2010).

#### Normalized mean distributions around putative RNase E sites and nascent and total RNA 3′ end peaks

For alignment of 3′ and 5′ end coverage relative to putative RNase E sites and RNA polymerase pauses, we focused on sites within protein-coding genes including adjacent UTRs (see definition of mRNA encoding genomic regions above). We defined symmetrical 60 nt windows around these sites and extract coverage information for all positions. For further analysis, only sites were considered that met our read cutoff criteria (0.5 reads per position in wildtype & Δ*pnp* for putative RNase E sites in Fig. 3, S3, 0.15 reads per position in polysome gradient samples for RNA polymerase pause sites, 0.15 reads per position in polysome gradient samples for RNA polymerase pause sites and RNase E sites in Fig. 4B). To be able to consider many different sites despite differences in (local) expression for analysis, coverage around each site was normalized by the mean coverage in the 60 nt window. We refer to this as relative coverage. A value of 1 resembled the region mean, everything above 1 was increased relative to the mean and everything between 0 and 1 decreased. For visualization, the composite mean of all relative coverages is plotted for each position centered around the sites of interest. In Fig. 3B, S3 the full 60 nt window is shown and in Fig. 4B we show the mean signal at the respective site with error bars as standard error of the mean. For visualization as heatmap, the relative coverage for each site was plotted ranked by the RNase E site Z-score (see peak calling for the data in (Clarke et al., 2014)). In Fig. S3D we plot the mean of the 3′ end adenylation percentage per position for the same sites as in Fig. 3B and S3B-C.

#### Normalized mean distributions across CDSs relative to the start codon

For alignment of 3′ end coverage relative to the start of CDSs, we focused on the 816 first CDSs in our mRNA encoding genomic regions. We added 100 nt at the start of each CDS to include 5′ UTRs in the coverage analysis. We filter CDSs based on length and read coverage (CDS > 150 nt, mean read coverage per 100 nt 5′UTR + CDS at least 0.01 in the lowest coverage dataset (fraction 1, 538 genes)). As described above for sites in a 60 nt window, we normalized coverage in each CDS by the mean coverage in the whole region. Next, we applied a running average smoothing per region within a 100 nt window. The smoothed relative coverage data were winsorized to the 95% quantile for each region. Finally, we computed the mean of the smoothed coverage at every position for −100 nt to 1900 nt across all CDSs that were at least as long as the respective position. Thus, more CDSs were included in the smaller distances to the start codon than towards later positions and the profile became noisier towards later positions (Fig. 4C).

### 3′ end adenylation of mature 3′ ends and RNase E sites

To investigate the global prevalence of 3′ end adenylation at mature 3′ ends and putative RNase E sites internal to protein-conding genes in wildtype, *Δpnp*, and *ΔpcnB*, we included all protein-coding genes in the analysis. Nucleotide positions that were within the mRNA encoding genomic regions, but not within CDSs, and showed 3′ end peaks with Z-scores greater 7 were considered mature 3′ ends (1341 (wildtype), 2003 (*Δpnp*), 905 (*ΔpcnB*) positions). The percentage of 3′ end adenylation for these sites was plotted as cumulative distribution for all three data sets in Fig. 3F with a minimum of 10 reads per mature 3′ end. Similar distributions were obtained for simple TUs only. We considered putative RNase E sites within mRNA encoding genomic regions and with 10 or more 3′ end reads (135 sites) for the quantification of 3′ end adenylation in Fig. 3F. To visualize the distribution of adenylation at and around putative RNase E sites (Fig. S3D), we included all putative RNase E sites that have at least 30 adenylated and non-adenylated reads in a 60 nt window (1423 sites). Hence, the average adenylation is given for the same sites as in Fig. 3B and S3B-C.

### Published data integration

Data from the following studies were remapped and integrated into the analysis described above:

- Term-seq from *E. coli* in LB at OD 0.5 (ERR2433552, ERR2433553, ERR2433554) and *B. subtilis* in LB OD 0.1-0.2 (ERR1232477, ERR1248362, ERR1248363) (Dar et al., 2016; Dar and Sorek, 2018).
- 5′ end-seq data rne-3071 ts (GSM1405878), rne wild-type (GSM1405877) (Clarke et al., 2014)
- Rend-seq data from *E. coli* MG1655 (GSM2500131) and *B. subtilis* 168 (GSM2500127)(Lalanne et al., 2018)
- Scripts used for read processing and mapping can be found on https://github.com/gwlilabmit/Herzel2022_RNAdemographics.

We used the reported mRNA half-lives obtained by τ-seq in (Moffitt et al., 2016) to compare to our Φ_end_ distribution (Fig. S3A). Only CDSs were considered with half-lives differing less than two-fold between τ-seq replicates.

Data analysis presented in this manuscript was done using bash scripts and software as cited above, python 2.7, matlab R2021b, Rstudio (Version 1.2.5042) and on the high-performance cluster LURIA maintained by the Biomicrocenter at MIT.

## Data access

The sequencing data from this study have been submitted to the NCBI Gene Expression Omnibus (GEO; http://www.ncbi.nlm.nih.gov/geo/) under accession number GSE189181. Scripts used for read processing and mapping can be found on github.com/gwlilabmit/Herzel2022_RNAdemographics.

## Acknowledgements

The authors thank Matthew H. Larson and Jason Peters for sharing the Flag-tagged RpoC strains, Tania A. Baker, Christopher B. Burge, Eliezer Calo, Joseph H. Davis, Alan D. Grossman, Rebecca L. Lamason and Michael T. Laub and their labs for sharing equipment and strains, Jean-Benoit Lalanne and Cassandra Schaening for providing python and matlab scripts for peak calling, Mirae Parker for assistance with the initial execution of the in-house RNA-seq protocol and manuscript comments, Matthew Z. Tien for the initial version of the in-house Northern blot protocol and explanation, and Michael R. Weber also for comments on the manuscript. This research was supported by the Helen Hay Whitney foundation (to LH) and the NSF Career Award MCB-1844668.

## Author contributions

LH and GWL conceived the study. JAS performed a P1 transduction to obtain replicate strains for *Δpnp*, measured the bacterial growth shown in Fig. S5A, performed polysome gradient centrifugation and Northern blot analysis of one wildtype and *Δpnp* replicate (Fig. S5B-D), and performed the +/- Kasugamycin treatment and Rend- and 3′ end-seq library preparation, data processing, mapping and expression quantification (Fig. 5D). CCY performed initial tests of different RNA extraction methods. LH performed all other experiments, analysis and prepared all the figures. LH wrote the first draft of the manuscript. GWL and JAS edited the manuscript.

## Supplemental Tables

**Supplemental Table 1:** Strains used in this study.

**Supplemental Table 2:** Oligonucleotides used in this study.

**Supplemental Table 3:** rRNA annotation used in this study.

**Supplemental Table 4:** Simple transcription unit annotation *E. coli*.

**Supplemental Table 5:** Simple transcription unit annotation *B. subtilis*.

**Supplemental Table 6:** Annotation of mRNA encoding regions *E. coli*.

**Supplemental Table 7:** Φ_end_ from 3′ end-seq.

**Supplemental Table 8:** Nascent RNA fraction.

**Supplemental Table 9:** RNase E-dependent 5′ ends.

## Figure legends

**Figure S1:**
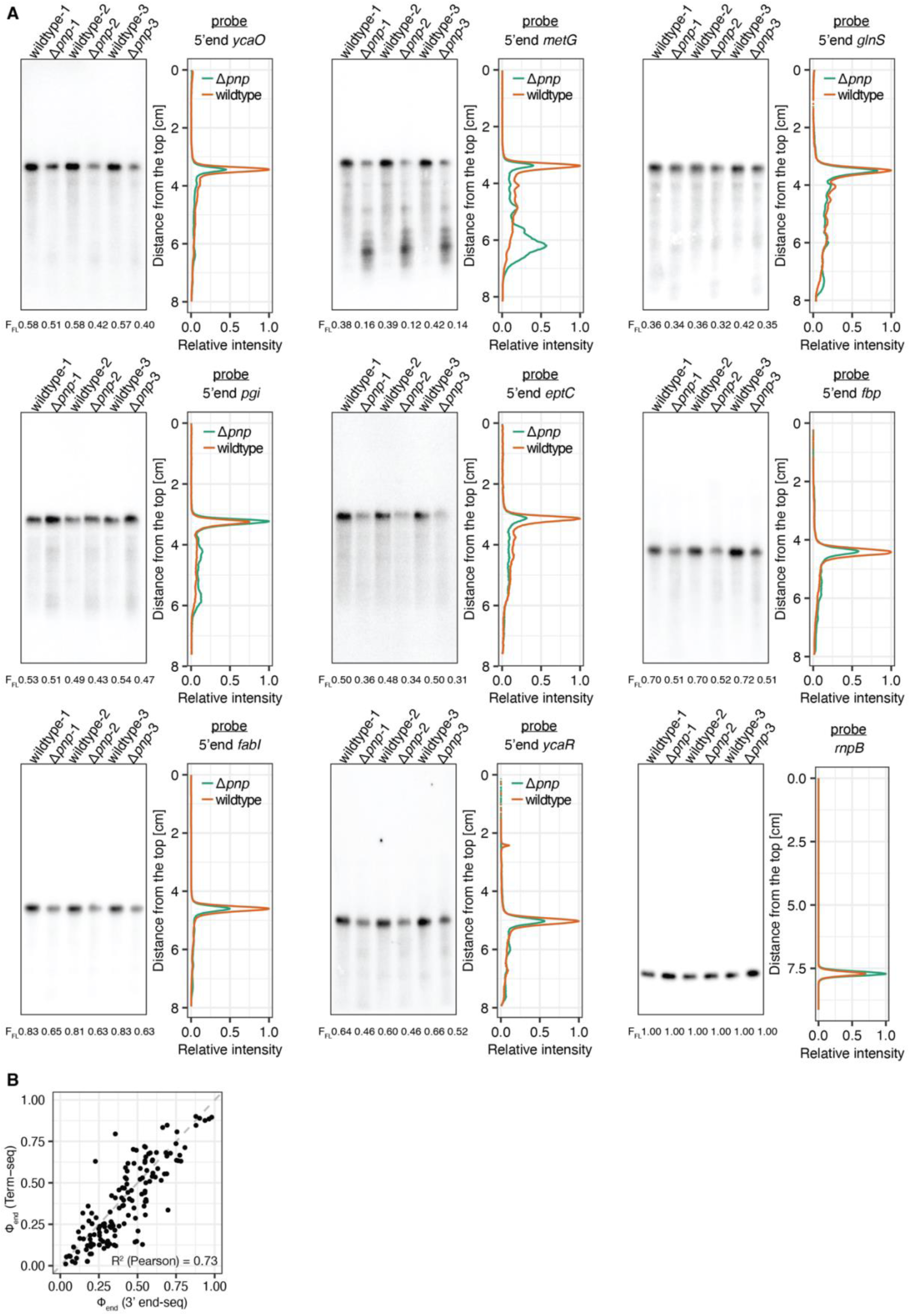
Measuring transcriptome integrity and heterogeneity by RNA-seq and Northern blot. (A) Northern blots and intensity profiles used for quantification of the full-length fraction in wildtype in Fig. 1F. 5 μg of total RNA from 3 biological replicates per strain. Probes bind in the 5′ region of TUs. Blotting against the stable RNA *rnpB* serves as loading control (last panel). (B) Reproducibility of Φ_end_ between different studies. Scatter plot shows Φ_end_ from simple TUs from this study and Term-seq (Dar and Sorek, 2018).

**Figure S2:**
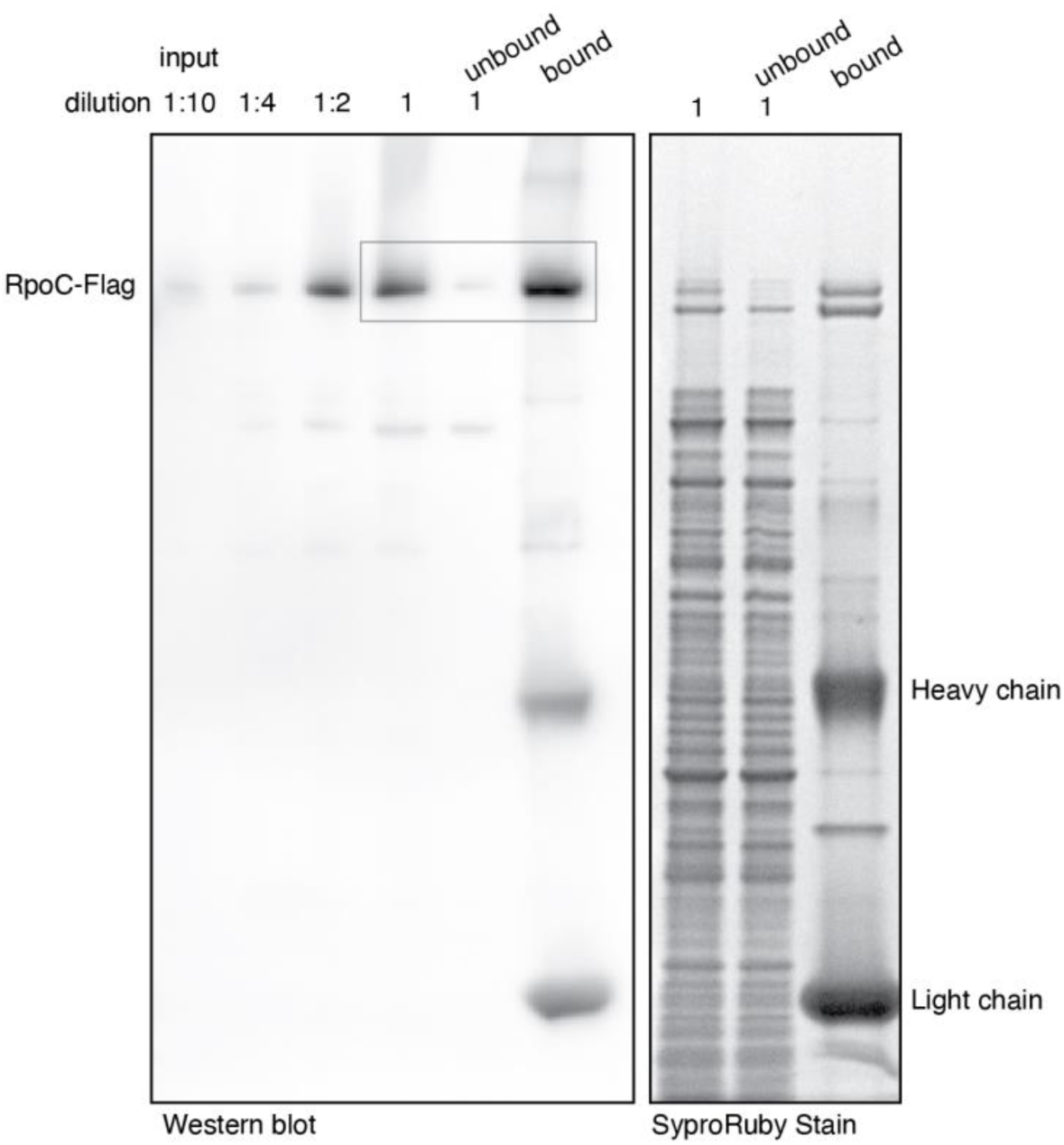
Probing for RNA polymerase enrichment. Full Western blot (left) probing for Flag-tagged RpoC and SyproRuby stain (right) after SDS-PAGE. The SyproRuby stain served as loading control and was prepared at the same time and with the same sample used in Western blot. Boxed area in the Western blot corresponds to the cutout shown in Fig. 2A. A dilution series was loaded for the input sample in the Western blot analysis to estimate the remaining RpoC after immunoprecipitation (10-25%).

**Figure S3:**
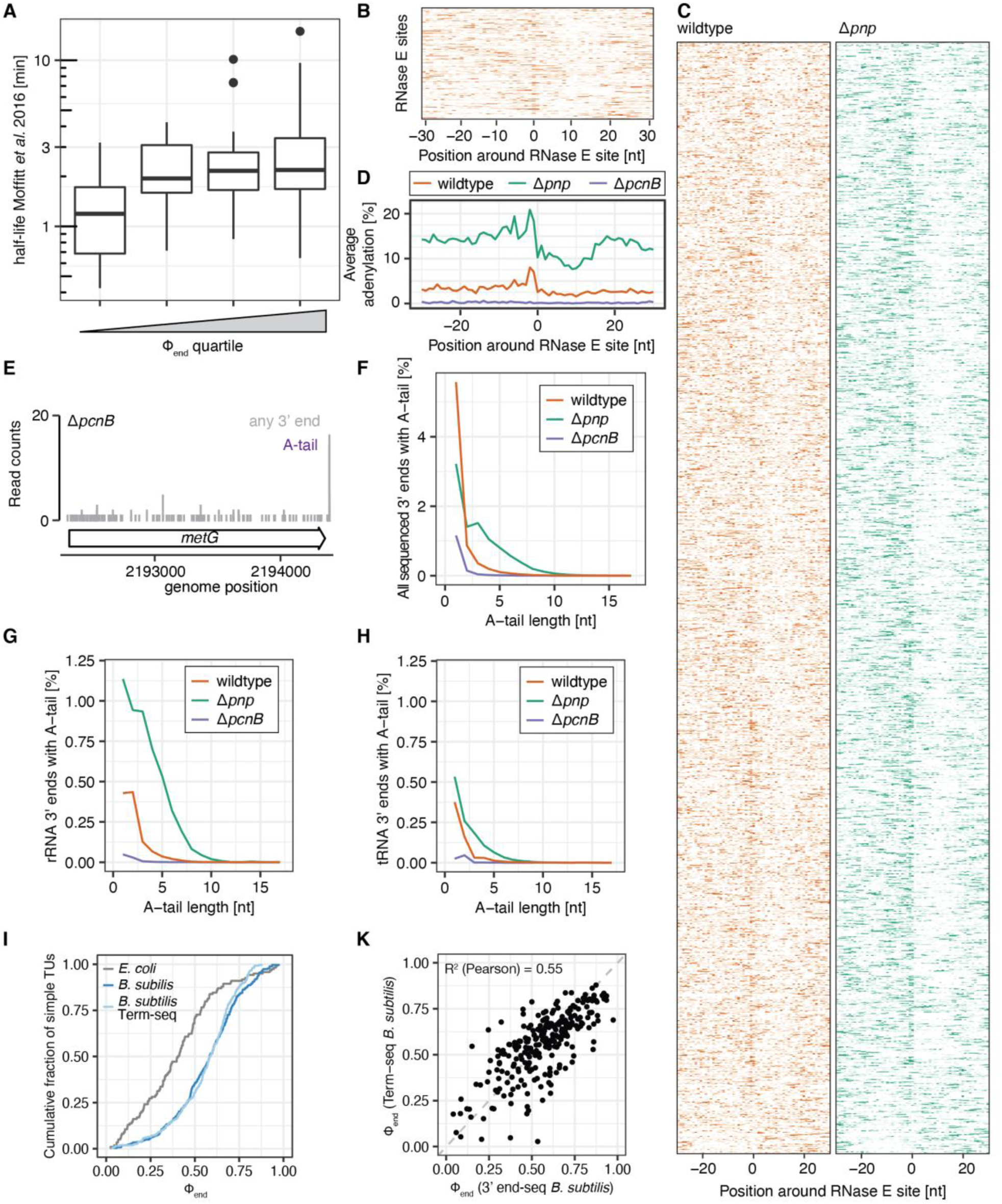
RNA decay signatures in mRNA, tRNA and rRNA. (A) Correlation between mRNA half-life and Φ_end_. Boxplot shows distributions of mRNA half-life for each quartiles of Φ_end._ Half-lives were determined by τ-seq (Moffitt et al., 2016). (B) Average wildtype 3′ end signals surrounding putative RNase E cleavage sites that have at least 30 reads in a 60-nt window. For each site, the read count for each position is normalized to the average read count in a symmetric 60-nt window. Normalized 3′ end read counts across individual RNase E sites in for wildtype. Each row represents data from each putative RNase E site. Color intensity represents the normalized 3′ end signal across the 60-nt window. Rows are sorted by descending Z-score at the putative cleavage site. The 200 sites with the highest number of reads in the 60-nt window are shown. (C) Full heatmaps complementing Fig. 3B and Fig. S3B. (D) Average percent adenylation surrounding the 1423 putative RNase E cleavage sites that have at least 30 reads in a 60-nt window for wildtype, *Δpnp*, and *ΔpcnB*. (E) Adenylation in the absence the PolyA polymerase PcnB. 3′ end profiles across *metG* are shown. (F-H) Percent of 3′ ends with A-tails of different lengths for (F) all mapped and unmapped reads, (G) in rRNA and (H) in tRNA for wildtype, *Δpnp* and *ΔpcnB*. As shown for mRNA in Fig. 3G, the average A-tail length is very short and increases upon deletion of the exonuclease PNPase (*Δpnp*). (I) Cumulative distributions of Φ_end_ for simple TUs in *E. coli* and *B. subtilis*. (K) Scatter plot correlating Φ_end_ from *B. subtilis* simple TUs from this study and Φ_end_ from a reanalysis of published Term-seq data (Dar et al., 2016).

**Figure S4:**
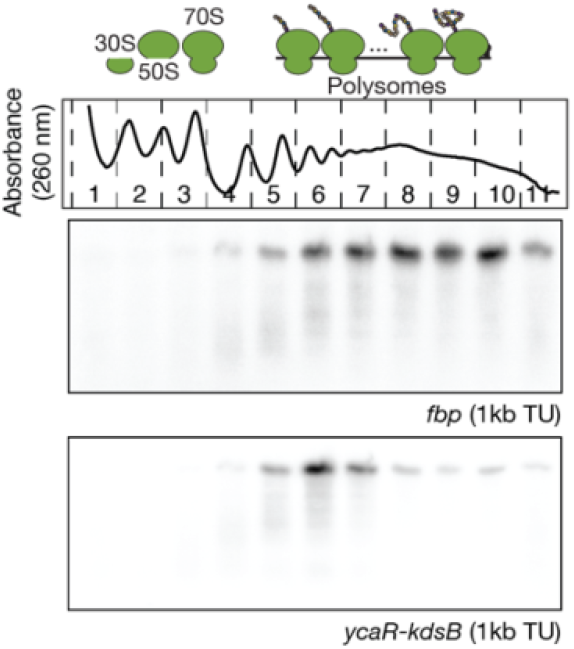
Additional Northern blot analyses for different TUs. Top: Absorbance profile across the sucrose gradient. Fraction numbers and the peaks that associate with 30S, 50S, 70S and polysomes are labeled. Bottom: Northern blot analyses of *fbp* and *ycaR-kdsB*, respectively. Northern blot probes hybridize to the 5′ region of the TU for *fbp* and *ycaR-kdsB*. Polysome gradient is same as in Fig. 4D and Fig. S5B.

**Figure S5:**
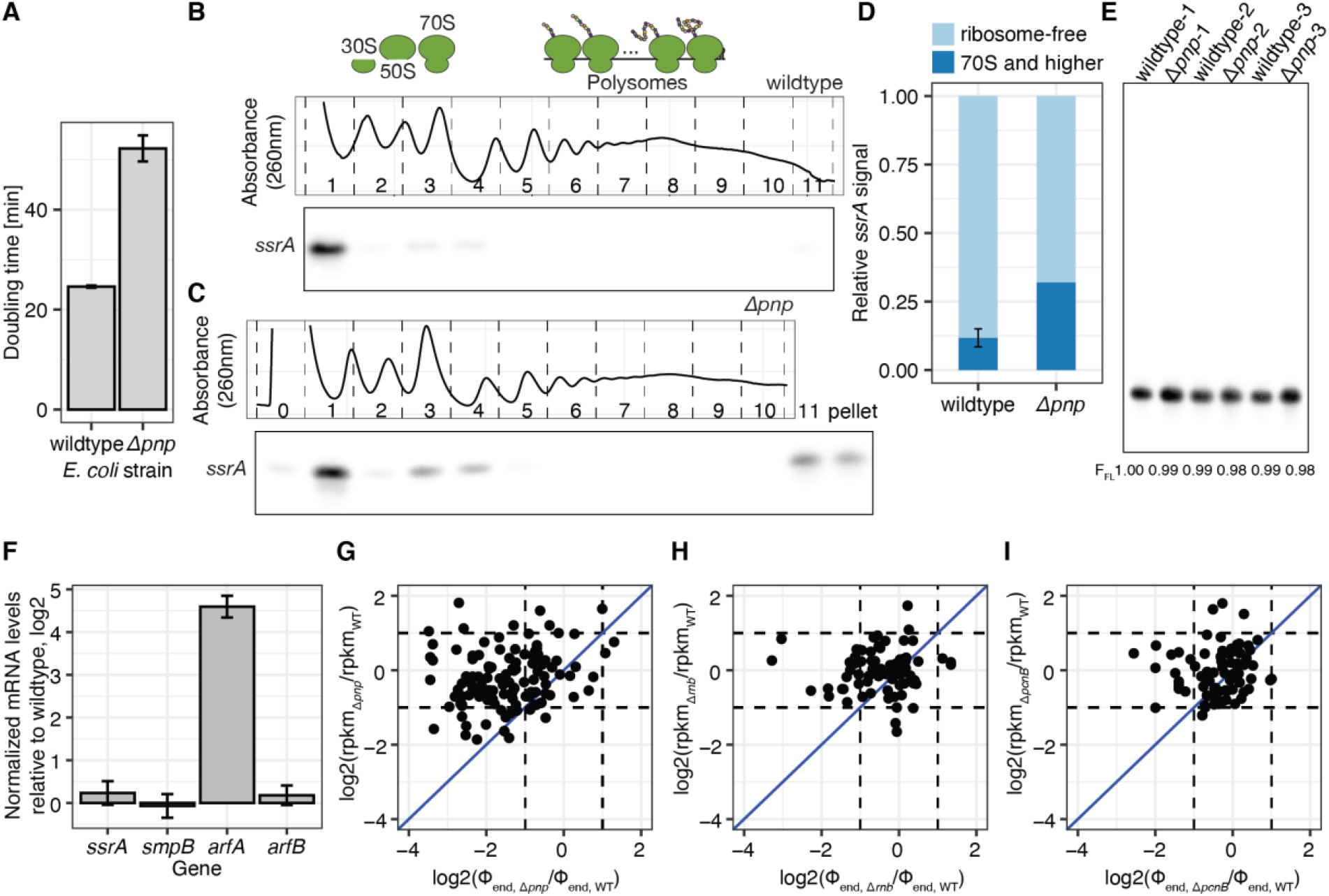
Higher engagement of *trans*-translation in *Δpnp* and transcriptome heterogeneity changes in deletions of RNA decay associated enzymes. (A) Doubling times for wildtype *E. coli* and *Δpnp*. Doubling times are measured in exponential phase in MOPS-Complete media with three independent biological replicates. Error bars correspond to standard deviation. (B) Northern blot profile of the RNA-component of the *trans*-translation machinery *ssrA* across polysomes for wildtype. (C) Similar to B, but for strain *Δpnp*. (D) Quantification of *ssrA* signal in ribosome-free (fraction 1,2 and 0-2, respectively in WT and *Δpnp*) and ribosome-associated (fraction >2) fractions. Error bar indicates standard deviation of 3 biological replicates in wildtype. (E) *ssrA* expression across different strains. Northern blot analysis for *ssrA* in total RNA shows that its expression level is not significantly different between wildtype *E. coli* and *Δpnp* (Student’s t-test, p>0.05). (F) mRNA expression of *trans-*translation system and alternative ribosome rescue factors. Relative expression between Δ*pnp* and wildtype is measured by RT-qPCR for 3 biological replicates. Error bars indicate standard deviation. (G-I) Fold-changes in Φ_end_ and rpkm values for simple TUs. Fold-change is calculated relative to wildtype for Δ*pnp* (replicate for Fig. 5C) (G), Δ*rnb* (H), and Δ*pcnB* (I).

